# Cumulative distribution functions: An alternative approach to examine the triggering of prepared motor actions in the StartReact effect

**DOI:** 10.1101/2020.04.23.056929

**Authors:** Aaron N. McInnes, Juan M. Castellote, Markus Kofler, Claire F. Honeycutt, Ottmar V. Lipp, Stephan Riek, James R. Tresilian, Welber Marinovic

**Affiliations:** School of Psychology, Curtin University, Perth, Australia; National School of Occupational Medicine, Carlos III Institute of Health, Madrid, Spain; Department of Neurology, Hochzirl Hospital, Zirl, Austria; School of Biological and Health Systems Engineering, Arizona State University, Tempe, AZ, United States of America; School of Human Movement and Nutrition Sciences, University of Queensland, Brisbane, Australia; Department of Psychology, University of Warwick, Coventry, United Kingdom

**Keywords:** motor control, muscle, reaction time, startle, StartReact effect, sternocleidomastoid

## Abstract

There has been much debate concerning whether startling sensory stimuli can activate a fast-neural pathway for movement triggering (StartReact) which is different from that of voluntary movements. Activity in sternocleidomastoid (SCM) electromyogram is suggested to indicate activation of this pathway. We evaluated whether SCM activity can accurately identify trials which may differ in their neurophysiological triggering and assessed the use of cumulative distribution functions (CDFs) of reaction time (RT) data to identify trials with the shortest RTs for analysis. Using recent datasets from the StartReact literature, we examined the relationship between RT and SCM activity. We categorised data into short/longer RT bins using CDFs and used linear mixed effects models to compare potential conclusions that can be drawn when categorising data on the basis of RT versus on the basis of SCM activity. The capacity of SCM to predict RT is task-specific, making it an unreliable indicator of distinct neurophysiological mechanisms. Classification of trials using CDFs is capable of capturing potential task- or muscle-related differences in triggering whilst avoiding the pitfalls of the traditional SCM activity based classification method. We conclude that SCM activity is not always evident on trials that show the early triggering of movements seen in the StartReact phenomenon. We further propose that a more comprehensive analysis of data may be achieved through the inclusion of CDF analyses. These findings have implications for future research investigating movement triggering as well as for potential therapeutic applications of StartReact.

## 1.0 Introduction

Large reductions of reaction time (RT) can be observed when an intense sensory stimulus is presented during movement preparation (Valls-Solé, Rothwell, Goulart, Cossu, & Muñoz, 1999), a phenomenon termed the StartReact effect. These observations of remarkably short RTs have led to the proposal that triggering mechanisms separate to those responsible for voluntary movements are activated by an intense sensory stimulus which is capable of producing a startle response (Carlsen, Dakin, Chua, & Franks, 2007; Carlsen, Maslovat, & Franks, 2012; J. Valls-Solé et al., 1999). That is, prepared movements may be released when a startling stimulus excites subcortical structures, bypassing the usual cortical circuits involved in voluntary motor control (Carlsen, Chua, Inglis, Sanderson, & Franks, 2004; Valls-Solé et al., 1999). Given the assumption that excitation of subcortical structures associated with the startle response can lead to the engagement of a distinct StartReact pathway for movement triggering, the presence of a startle response has traditionally been used to differentiate the StartReact effect from other phenomena that can cause (usually less extensive) reductions in RT, such as the well-documented stimulus intensity and accessory stimulus effects (Bernstein, Clark, Edelstein, & Grant, 1969; Pieron, 1914; Pins & Bonnet, 1996). Thus, motor responses have typically been defined as StartReact movements on the basis of activity in surface electromyography (EMG) of the sternocleidomastoid (SCM) muscle, which is said to indicate startle (Carlsen et al., 2007). When no SCM activity is recorded in a trial, it is assumed that the specific mechanism responsible for the StartReact effect was not activated, and the less dramatic reductions of response time that are typically observed are attributed to stimulus intensity and/or accessory stimulus effects through the pathway used for volitional motor control (Carlsen, Maslovat, Lam, Chua, & Franks, 2011; Kohfield, 1971).

While, on average, movements in the presence of SCM activity usually occur with shorter RTs than those in absence of SCM activity, it has not been unequivocally demonstrated that observation of a startle response is a necessary condition for the vast reductions of RT which are indicative of the StartReact effect (Marinovic & Tresilian, 2016). For example, surface SCM activity is not always present when eliciting movements with latencies short enough to be indicative of a StartReact effect. The impaired reliability of using SCM as a marker of neurophysiological circuitry is further demonstrated by the finding that SCM activity can be reduced with pre-pulse inhibition without modifying RT shortening in the StartReact, and unlike startle, the StartReact effect does not appear to be prone to habituation (Castellote, Kofler, Mayr, & Saltuari, 2017; Leow et al., 2018; Marinovic & Tresilian, 2016; Valldeoriola et al., 1998; J. Valls-Solé, Kofler, Kumru, Castellote, & Sanegre, 2005). As such, even if activity associated with the intense stimulus reaches startle-related circuits, this may not always be indicated by SCM activity. Therefore, making inferences about the circuitry used for fast movement triggering based on surface SCM activity may be rather unreliable. Furthermore, the available data do not preclude cortical involvement in the StartReact effect. As an alternative view to this triggering through subcortical areas, the shortening of RT seen in the StartReact effect may be a product of an enhancement of voluntary motor pathways via an engagement of a more wide-spread cortical-subcortical network when an intense sensory stimulus is presented (see Marinovic & Tresilian, 2016). The difficulties in determining neurophysiological mechanisms underlying the early triggering of motor responses using the presence or absence of SCM activity has been outlined previously (Dean & Baker, 2017; Leow et al., 2018; Marinovic & Tresilian, 2016; McInnes, Corti, Tresilian, Lipp, & Marinovic, 2020) and it seems SCM activity can be an unreliable indicator of distinct mechanisms that can be activated by intense sensory stimuli. Rather, determination of the presence of a specific StartReact mechanism may be more feasible when trials are separated based on their response latency (Leow et al., 2018; McInnes et al., 2020).

Here, we evaluate the utility of separating RT trials on the basis of SCM activity to investigate mechanisms underlying the StartReact phenomenon and further examine an alternative approach, cumulative distribution functions (CDFs) that separate trials based on response latency (Leow et al., 2018; McInnes et al., 2020). CDFs allow an examination of how trials with the fastest RTs differ from those with slower RTs which would be considered unrepresentative of the StartReact effect, whilst avoiding the pitfalls of relying on SCM activity as an indicator of StartReact mechanisms which have been outlined previously (Marinovic & Tresilian, 2016). We re-analysed data from seven studies (Castellote & Kofler, 2018; Honeycutt, Kharouta, & Perreault, 2013; Honeycutt, Tresch, & Perreault, 2014; Marinovic, de Rugy, Riek, & Tresilian, 2014; Marinovic, Milford, Carroll, & Riek, 2015; Ossanna, Zong, Ravichandran, & Honeycutt, 2019; Tresch, Perreault, & Honeycutt, 2014) which have investigated differences in response times across trials in the presence and absence of SCM activity. We used our method to evaluate the utility of separating trials on the basis of SCM activity by examining the distribution of SCM activity across the spectrum of RTs and evaluating the relationship between RT and the presence of SCM activity. We further analysed these datasets in order to define a common method of separating trials on the basis of response latency. Lastly, we used our method of trial categorisation to evaluate the hypothesis that separate mechanisms contribute to StartReact and voluntary movements.

## 2.0 Methods

Data comparing responses which occur in the presence and absence of SCM activity were provided from the authors of seven studies reported in recent literature and subject to statistical analyses. Note that the Tresch et al. (2014) dataset includes data collected from participants with stroke which were reported separately in Honeycutt et al. (2014). For the sake of brevity, we have limited the report within the main body to the analysis of a single dataset provided by Castellote and Kofler (2018). This task recorded EMGs from the biceps brachii (BB) in a flex-only task, first dorsal interosseous (FDI) in a pinch-only task, and both BB and FDI in a combined pinch-flex task. Extended analyses for the individual datasets from the remaining studies, which differed in tasks used and muscles from which EMGs were recorded in addition to SCM (summarised in Table 1), are reported in the appendices of this report.

**Table 1.** Overview of studies included in analyses.

| *Authors (year)* | *N* | *Task* | *Muscles recorded* |
| --- | --- | --- | --- |
| *Castellote & Kofler*<br>*(2018)* | 11 | Elbow flexion | Biceps brachii (BB) |
|  |  | Finger pinch | First dorsal interosseous (FDI) |
|  |  | Combined pinch-flex | BB and FDI |
| *Honeycutt et al.*<br>*(2013)* | 10 | Finger abduction | FDI |
|  |  | Grasp |  |
| *Marinovic et al.*<br>*(2014)* | 7 | Lip press | Orbicularis oris (OO) |
|  |  | Button press with thumb | Abductor pollicis brevis (APB) |
| *Marinovic et al.*<br>*(2015)* | 10 | Arm supination | BB |
| *Ossanna et al. (2019)* | 10 | Five-direction arm reaching | Anterior deltoid (AD) |
|  |  |  | BB |
|  |  |  | Brachioradialis (Br)<br>Posterior deltoid (PD)<br>Pectoralis (Pe)<br>Triceps brachii (TB) |
| *Honeycutt et al.*<br>(2014); *Tresch et al.*<br>(2014) | 34 | Hand flexion, extension | Extensor digitorum<br>communis (EDC)<br>Flexor digitorum superficialis<br>(FDS)<br>FDI |

All analyses were conducted using R software (*v3.6.0*; R Core Team, 2019). StartReact experiments typically employ “control” trials in which the participant performs a predetermined movement in response to an imperative stimulus (IS). In a subset of trials, an intense sensory stimulus (probe) is delivered in addition to the IS. Data used for our analyses of each individual dataset were limited to (premotor; time to EMG onset) RTs in probe trials for which an intense sensory stimulus was delivered (i.e. control trials were removed). Movements made in response to probes for which SCM activity was recorded are defined as SCM+ responses. Responses not accompanied by SCM activity were defined as SCM− responses. If the responses of the target muscles in a given task that occur after an intense stimulus differ in terms of neurophysiological pathways, i.e. are either short latency SCM+ movements or longer latency SCM− movements, then RTs from those target muscles should fit a bimodal distribution. Alternatively, if a common mechanism underlies both SCM+ and SCM− movements, the data should fit a unimodal distribution. Data were separated for each task type and/or muscle type and we tested for the modality of each distribution with Hartigan’s (1985) dip test, using the *dip.test* function from the *diptest* package (*v0.75*). Due to the skewness commonly observed in RT data, we conducted a natural logarithmic transformation of all data for each movement type to assess whether skewness had any significant impact on the results of the dip tests (Whelan, 2008).

For all movements within each experiment, we calculated each participant’s median RT for SCM+ and SCM− responses. We conducted paired sample t-tests on these median values to examine the difference in RT between SCM+ and SCM− trials. This test allowed us to examine what muscles or movement types are identified as being amenable to StartReact in accordance with the SCM based method used to categorise responses to the intense probe stimulus. These results were later used for comparison with our analyses using the classification of responses on the basis of response latencies via CDFs.

For each individual participant, CDFs were calculated for the response time data of all trials in which an intense probe was delivered for each movement recorded, using the *quantile* function (Hyndman & Fan, 1996) from the *stats* package (*v3.6.0*). Quantiles were calculated for each participant’s RTs at the 5^th^, 15^th^, 25^th^, 35^th^, 45^th^, 55^th^, 65^th^, 75^th^, 85^th^, and 95^th^ percentiles of RT. We then calculated the mean RT across subjects for each of the quantiles within the CDFs (Ratcliff, 1979), giving ten values which represent the average response times of participants at each percentile for all CDFs we conducted. We further calculated the mean of our subject medians of SCM+ and SCM− responses to determine the mean SCM+ and SCM− latencies. Once these were calculated, these average SCM+ and SCM− latencies were used to estimate the latencies of responses that may differ in their triggering mechanisms and compared these to our calculated quantiles. Therefore, given SCM+ and SCM− trials have been assumed to differ in their triggering mechanisms, for a given movement type within each experiment, the mean percentile closest in terms of RT to the mean SCM+ trial latency across participants was deemed the SCM+ percentile. Similarly, the mean percentile latency that was closest to the mean latency of SCM− trials for a given movement type was deemed the SCM− percentile. These percentiles allowed us to approximate the short and long RTs that may occur as a product of the potentially different neurophysiological pathways contributing to RTs in response to the intense probe stimulus.

Once percentiles approximating responses with and without SCM activity were calculated, we used these percentiles to group data into “startle” and “non-startle” categories (see Figure 1). If distinct mechanisms are activated for StartReact versus voluntary movements, splitting trials on the basis of latency should separate those movements which are thought of as being distinct, with trials at the shortest latencies representing the StartReact triggered movements and those at the longer latencies representing voluntarily triggered movements. Trials were placed into the startle category if their RT was equal to or shorter than the SCM+ percentile latency that was calculated for a given movement type within an experiment. Similarly, trials were placed into the non-startle category if their RT was equal to or longer than the SCM− percentile latency for that movement/muscle. We then calculated the percentage of trials within each category that occurred with SCM activity. This was calculated as 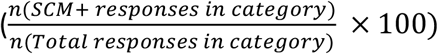. If SCM activity is a critical criterion for the considerable reductions of RT in the StartReact effect, then SCM+ responses should primarily occur in the startle category and SCM− responses should primarily occur in the non-startle category. To test this, we conducted a series of Bayesian tests of association using the *contingencyTableBF* function from the *BayesFactor* package (*v0.9.12*), with the joint multinomial sampling method (Albert, 1997; Gunel & Dickey, 1974; Morey et al., 2018). This test assesses the degree to which the data provide evidence for the dependence of SCM activity (SCM+/SCM−) on startle categorisation (startle/non-startle). If the presence of SCM activity does indeed depend on startle or non-startle categorisation – that is, SCM activity is predominantly found for responses in the startle category – this test would provide decisive evidence against the null hypothesis. This result would provide support for the use of SCM activity as an indicator of the activation of a fast-neurophysiological pathway. If, however, SCM+ responses are distributed across both startle and non-startle categories, and these variables are independent of one another, we expect to observe weaker evidence to support their dependence. BF_10_ values are reported, which describe the degree to which the data provide evidence against the null hypothesis.

**Figure 1.**
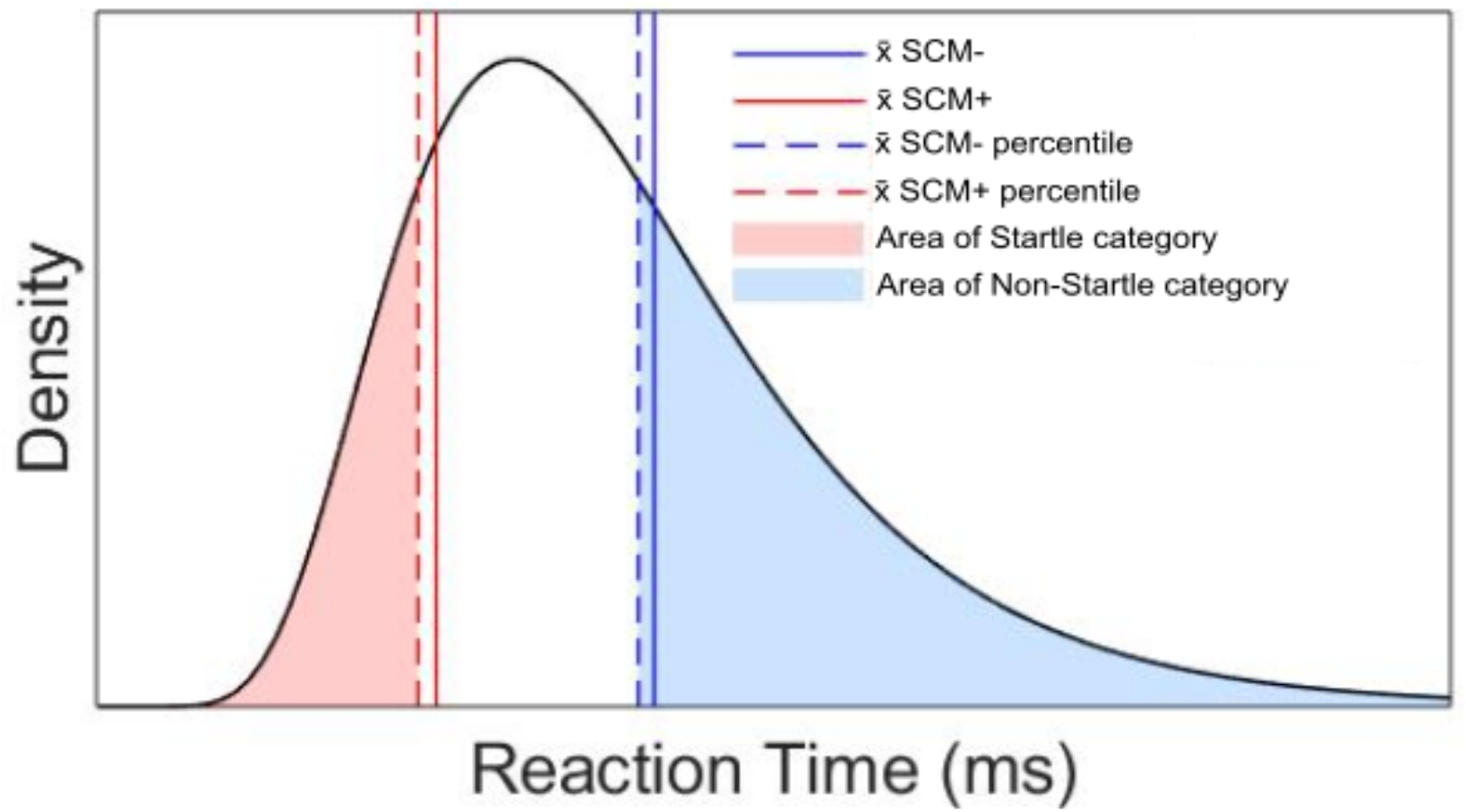
Illustration of the method used to categorise trials. Mean latencies of SCM+ and SCM− responses were used to determine the equivalent SCM+ and SCM− percentiles. Responses were categorised into the startle category if their latency was equal to or shorter than the mean latency of the SCM+ percentile. Responses were categorised into the non-startle category if their latency was equal to or longer than the mean latency of the SCM− percentile. Note this is a modelled exemplary RT dataset with x and y values not to scale.

We further used our percentiles across all data analysed to determine a common percentile ranking which may be used to categorise responses across all data sets. Across all tasks and muscles over all datasets analysed, the 45^th^ percentile was the latest percentile that was approximated to SCM+ responses, and the 55^th^ percentile was the earliest percentile approximated to SCM− responses. For each movement type in each experiment, responses at the 45^th^ percentile or earlier were therefore determined to be equivalent to (fast onset) SCM+ responses, and response at the 55^th^ percentile or later were determined to be equivalent to (slower onset) SCM− responses.

With our categorised data, we conducted a linear mixed-effects model with Kenward-Roger approximation for degrees of freedom using the *lmer* function (*lmerTest* package; *v2.0-36*; Kuznetsova et al., 2017) on the Castellote and Kofler (2018) data as a representative dataset. Percentile categorisation (fast onset/slower onset) and task type were set as fixed-factors in the model and participants were set as a random factor. To test whether distinct triggering mechanisms exist for StartReact versus voluntarily initiated responses, we examined the interactions of percentile and task/muscle type to assess whether the shortening of RT by the probe stimulus differs between movement types which likely have distinct connectivity to different brain regions. Post hoc analyses were conducted using the *emmeans* function (*emmeans* package; *v1.3.4;* Lenth, 2019) using the Tukey correction for multiple comparisons.

In order to encourage future use of CDFs when investigating triggering mechanisms in the StartReact effect, we have provided an R script which runs all analyses used in this report on a simulated dataset. The code can be obtained at http://doi.org/10.5281/zenodo.3760340. The data from the studies analysed in this report have been published elsewhere and may be obtained at the request of the original authors.

## 3.0 Results

### 3.1 Unimodality versus bimodality of data

Hartigan’s (1985) dip test failed to reject the null hypothesis of unimodality for the elbow flexion (flex-only; *p* = .715), the finger pinch (pinch-only; *p* = .095), the combined task BB latency (BB pinch-flex; *p* = .093), or the combined task FDI latency (FDI pinch-flex; *p* = .277) reported by Castellote and Kofler (2018). This suggests all tasks analysed produced a unimodal distribution of data. Extended analyses of the remaining datasets are presented in Appendix A. The analysis of the logarithmically transformed data was consistent with that of the original data and as such, we have reported the analyses of untransformed data.

### 3.2 Differences between SCM+ and SCM− trials

Paired sample t-tests of the difference between each subject’s median SCM+ and SCM− trial latencies for all movement types in the representative dataset indicated a significant difference in RT between SCM+ and SCM− responses in BB for the flex-only task (mean difference = −30.2 ms, CI = −38, −22.4), in BB for the combined pinch-flex task (mean difference = −53.5 ms, CI = −66.3, −40.7), in FDI for the combined pinch-flex task (mean difference = −54.6 ms, CI = −67.7, −41.6), but not in FDI for the pinch-only task (mean difference = −19 ms, CI = −38.5, 0.5). Extended analyses can be found in Appendix B.

### 3.3 Determining SCM+ and SCM− percentiles

For all tasks analysed, we have indicated the equivalent SCM+ and SCM− percentiles in Table 2. The percentage of responses within each category after splitting the data into startle and non-startle categories (see Figure 1) are also presented in Table 2. The CDFs calculated for the Castellote and Kofler (2018) data are plotted along with the mean latency of SCM+ and SCM− responses in Figure 2, and the distribution of SCM+ responses within the startle and non-startle categories can further be seen in Figure 3. Extended analyses are presented in Appendix C.

**Table 2.**
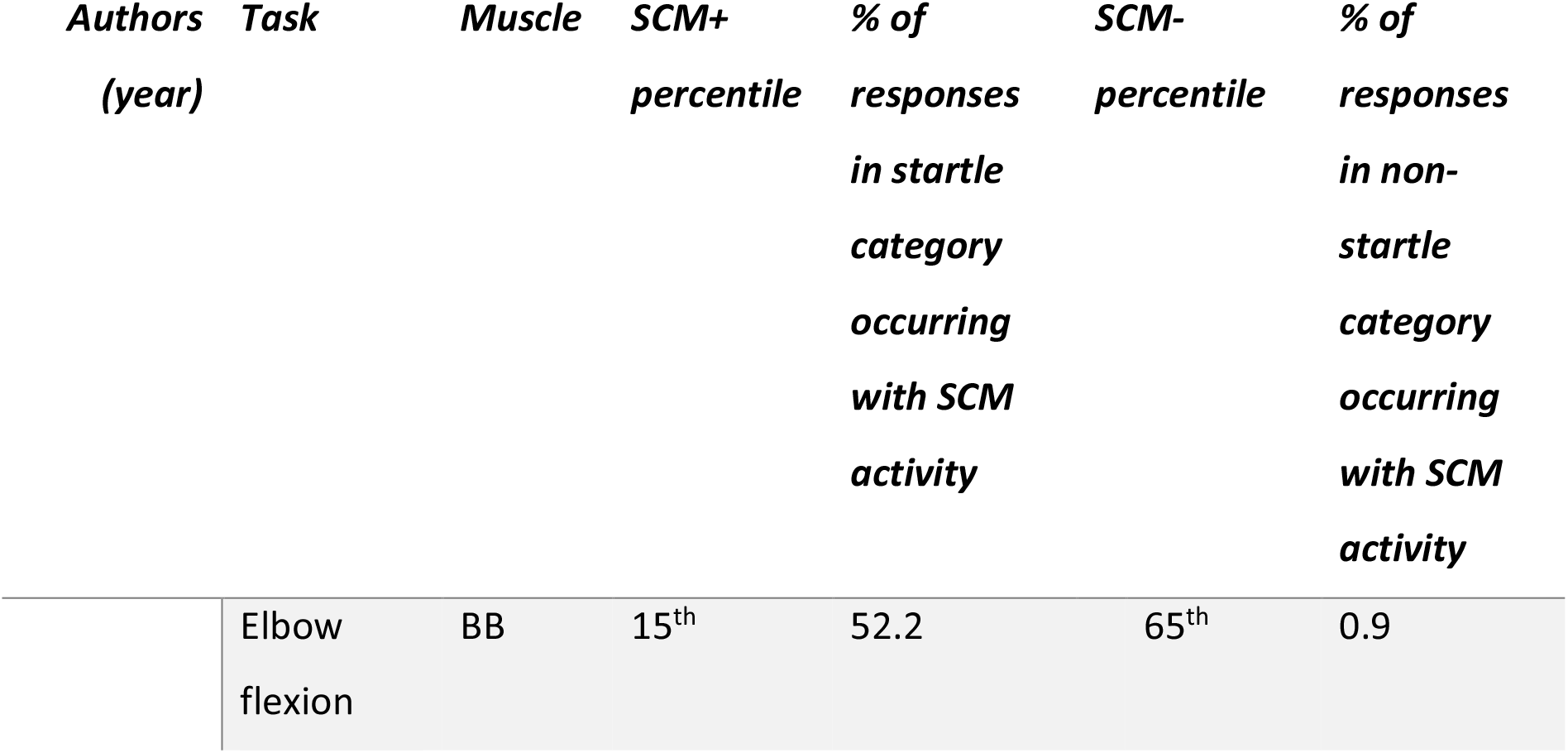
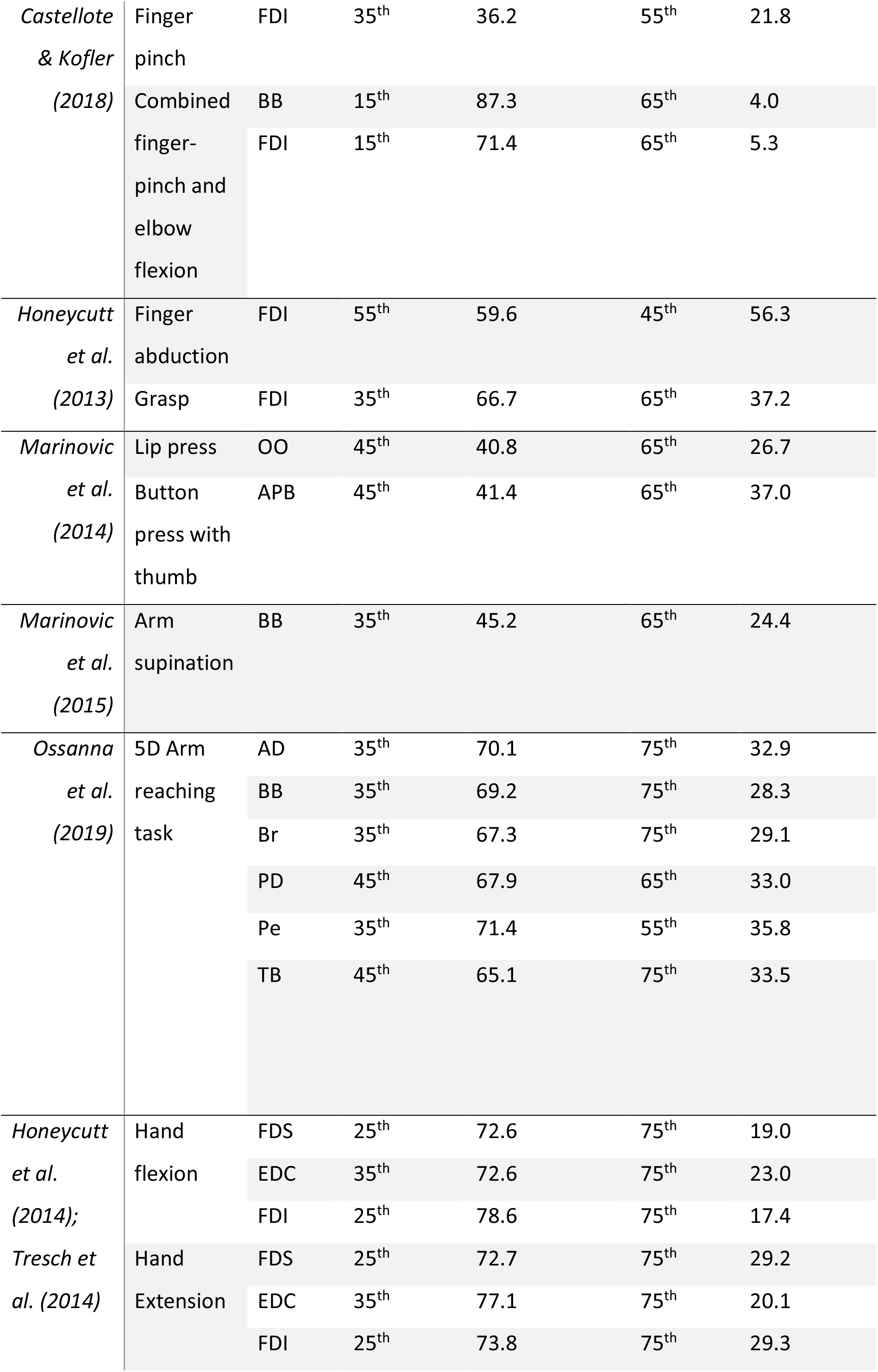
Overview of percentiles closest matching the mean latency of SCM+ and SCM− responses, along with the percentage of responses within the SCM+ and SCM− categories which occurred with SCM activity for each muscle and task analysed.

**Figure 2.**
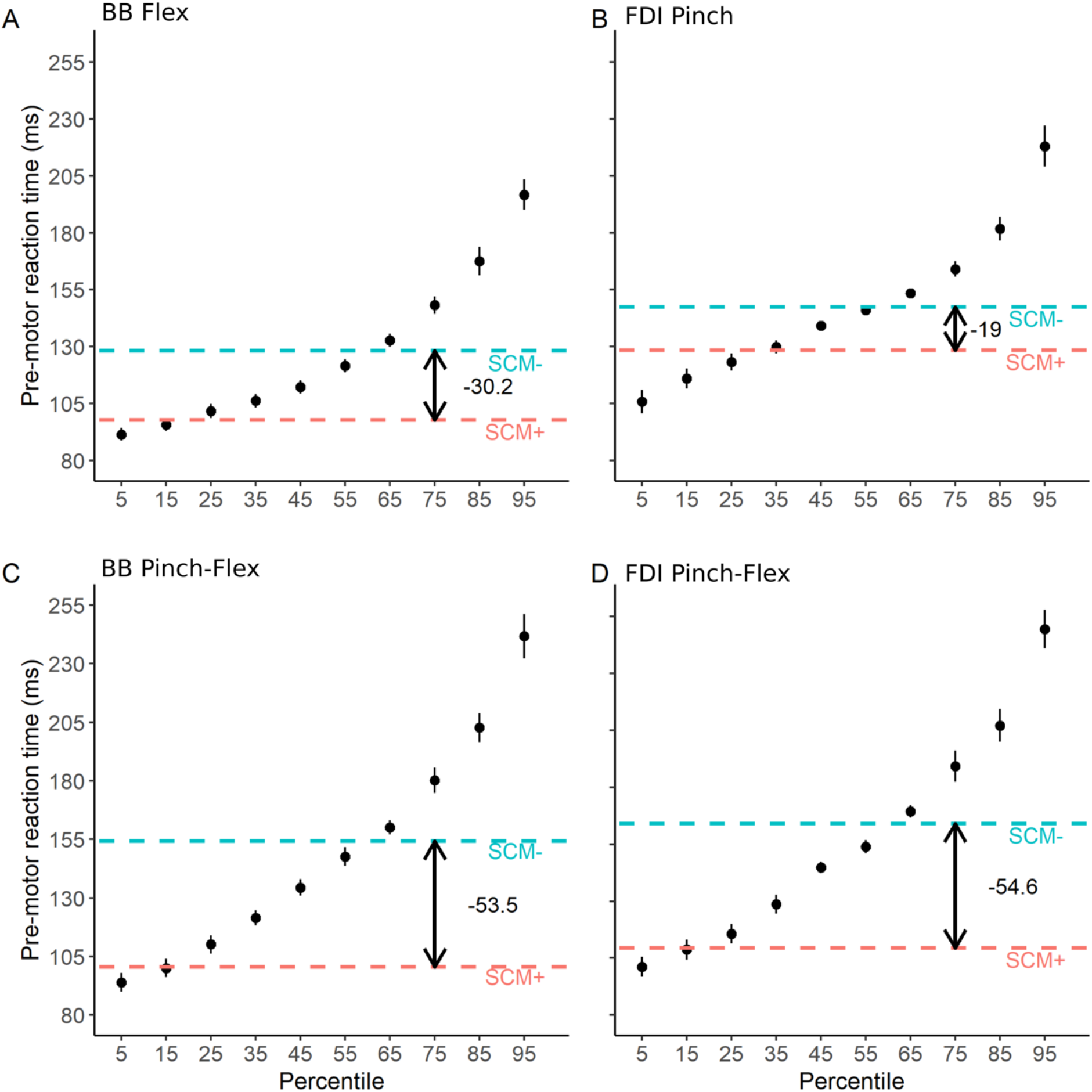
**Cumulative distribution function of Castellote and Kofler’s (2018) data.** A). Biceps brachii (BB) latency for flexion task B). First dorsal interosseous (FDI) latency for pinch task C). BB latency in combined task D). FDI latency in combined task. The mean latency of responses in the presence (SCM+) and absence (SCM−) of sternocleidomastoid activity are shown by the dotted lines. Error bars represent standard error of the mean.

**Figure 3.**
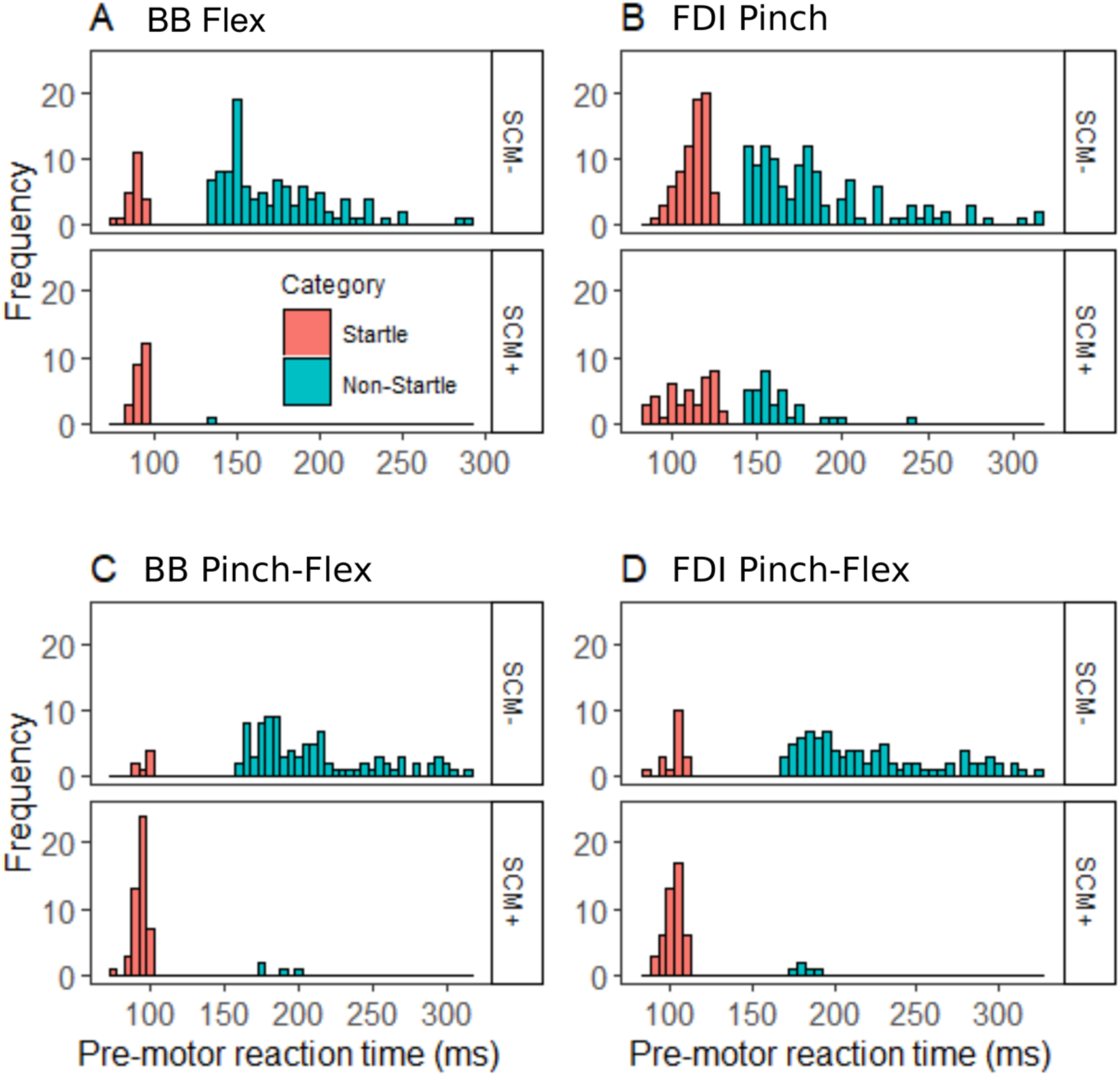
**Histogram displaying response times of SCM+ and SCM− responses across startle and non-startle percentile categories for Castellote and Kofler’s (2018) data.** A.) Biceps brachii (BB) latency in flex-only task. B). First dorsal interosseous (FDI) latency in pinch-only task. C). BB latency in combined pinch-flex task. D). FDI latency in combined pinch-flex task.

### 3.4 Presence of sternocleidomastoid activity in shorter and longer latency reaction times

Given some movement types showed a large proportion (max = 56.3%; see Table 2) of trials in the non-startle categorisation of RT which occurred with SCM activity, we conducted a Bayesian test of association (Albert, 1997) to examine whether the presence of SCM activity differs across our startle and non-startle RT categories. The analysis of Castellote and Kofler’s (2018) data resulted in BF = 2.5 × 10^12^ for BB in the flex-only task, BF = 6 for FDI in the pinch-only task, BF = 2 × 10^25^ for BB in the combined pinch-flex task, and BF = 8.2 × 10^16^ for FDI latency in the combined pinch-flex task. BFs > 100 indicate decisive evidence against the null hypothesis (Jeffreys, 1961) and as such, the flex-only, BB pinch-flex, and FDI pinch-flex tasks show decisive evidence for the dependence of percentile categorisation on SCM activity. These results indicate FDI latency in the pinch-only task is the only task within the dataset for which decisive evidence for the percentile-SCM dependence failed to be found. Extended analyses are shown in Appendix D.

### 3.5 Examining triggering mechanisms via faster onset and slower onset categorisation

Our analyses indicated that SCM activity does not always co-occur with shortened RT and also suggested that this relationship may be task-dependent. That is, for some tasks, a significant proportion of SCM+ responses are not only found in the startle category, but also within the non-startle category of RTs which approximates the longer latency SCM− RTs. Therefore, we examined an alternative approach to investigate triggering mechanisms of responses via intense sensory stimuli: categorisation via percentiles of RT. With the Castellote and Kofler (2018) dataset as a representative example, we conducted a linear mixed-effects model analysis to examine the appropriateness of this method of distinguishing responses at long (>= 55^th^ percentile) and short (<= 45^th^ percentile) latencies (see Figure 4). As expected, the main effect of percentile categorisation (fast onset/slower onset) was statistically significant, F_(1, 422)_ = 533.67, *p* < .001. More importantly, the interaction of percentile categorisation with muscle type (BB/FDI) was not statistically significant, F_(1, 422)_ = 0.05, *p* = .814, however, the interaction of percentile categorisation with task type (combined/single) was found to be statistically significant, F_(1, 422)_ = 18.63, *p* < .001.

**Figure 4.**
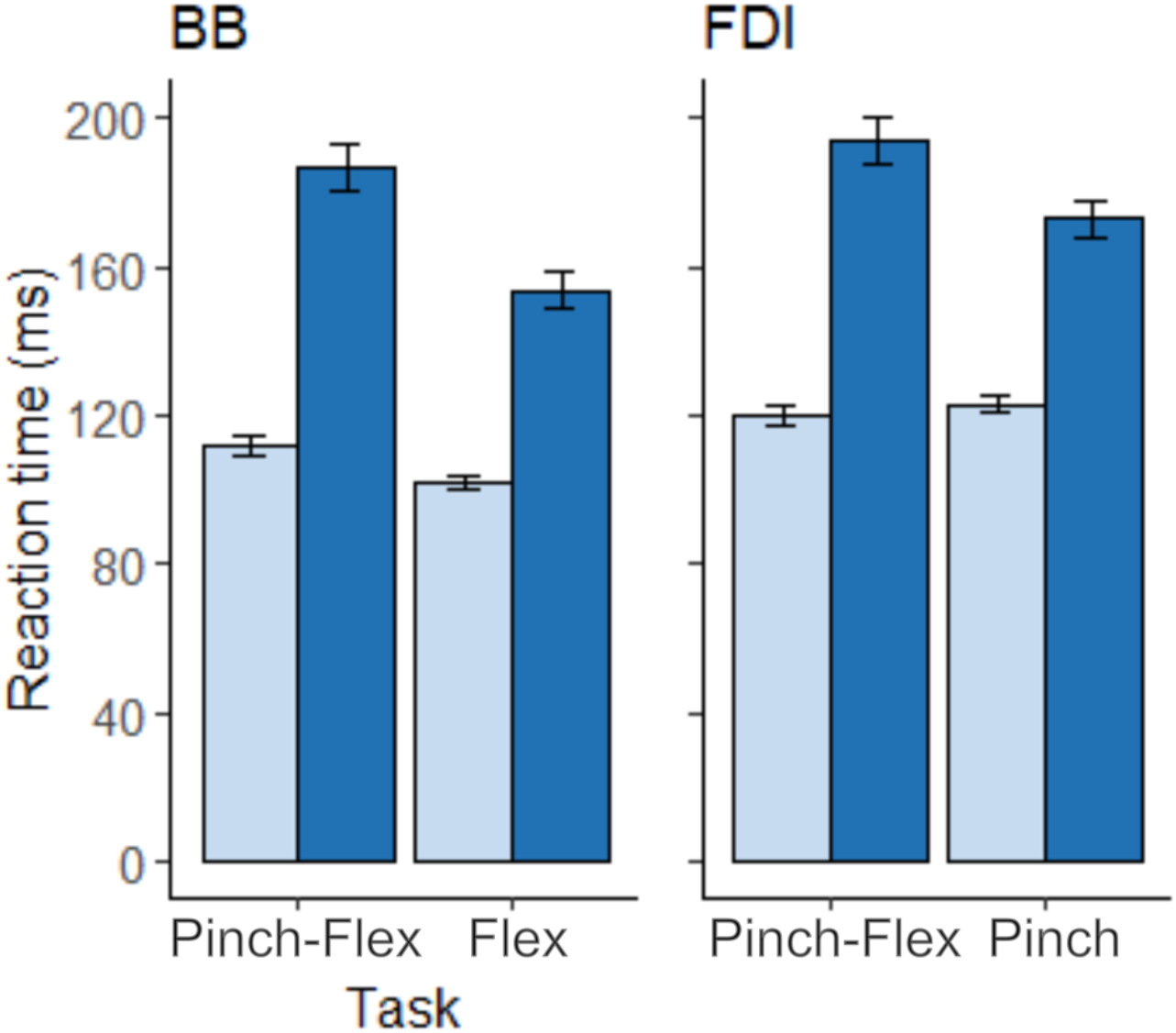
**Mean reaction time across subjects for all tasks reported by Castellote and Kofler (2018).** The data have been categorised into fast onset and slower onset response times. A). Biceps brachii (BB) latencies. B). First dorsal interosseous (FDI) latencies. Error bars represent standard error of the mean.

If separate mechanisms contribute to the fastest RTs – as a result of a modulated effect of the probe stimulus between muscles or tasks which differ in their neurophysiological contributions, then differences in RT should be observed between muscles or tasks in the fast-onset percentiles. Therefore, we ran a linear mixed model on the fast onset data to test the hypothesis that differences across tasks/muscles may be observed in trials at the shortest RTs. Our analysis found a statistically significant interaction of task type with muscle type, F_(1, 206)_ = 10.9, *p* = .001. Post-hoc analysis indicated a significant difference in RT between BB latency in the combined pinch-flex task and BB latency in the flex-only task, *p* = .002. This difference was not significant between FDI in the combined pinch-flex task and FDI in the pinch-only task, *p* = .725. The results of these analyses using our categorisation method via RT are consistent with those in the original report (Castellote & Kofler, 2018). Extended analyses can be seen in Appendix E.

We further analysed this dataset using the traditional classification of responses on the basis of SCM activity in order to compare the two methods of categorisation, and this produced similar results. Correspondingly with the previous main effect of percentile categorisation, the main effect of SCM activity (SCM+/SCM−) using the traditional method was statistically significant, F_(1, 1263.9)_ = 252.53, *p* < .001. Furthermore, in line with the observed statistically significant interaction of percentile categorisation with task type in our previous analysis, the interaction of SCM activity with task type was statistically significant, F_(1, 1260.8)_ = 25.49, *p* < .001, but the interaction of SCM activity with muscle type was not, F_(1, 1256.2)_ = 2.66, *p* = .103. Further examination of the SCM+ data showed an interaction of task type with muscle type, F_(1, 352.6)_ = 25.93, *p* < .001, consistent with that of the fast-onset data. However, post hoc tests indicated RTs of FDI in the pinch-only data (*M* = 132.95 ms, *SD* = 23.87) were significantly longer than those for BB in the flex-only data (*M* = 103.87 ms, *SD* = 17.66; *p* < .001), BB in the combined pinch-flex data (*M* = 108.56 ms, *SD* = 24.56; *p* < .001), and FDI in the combined pinch-flex data (*M* = 117.16 ms, *SD* = 24.47; *p* < .001). Importantly, these differences may be explained by our previous finding that for FDI latency in the pinch-only task there were a larger percentage (21.8%) of trials in the longer-latency percentiles of RT which occurred with SCM activity in comparison to the other tasks (see Table 2). Potentially these SCM+ trials at longer RTs would have had an impact on average latencies of FDI in the pinch-only data and as such, result in the significantly longer RTs for FDI in the pinch-only task as compared to the other tasks when categorising via SCM. This may lead to alternative interpretations of the data when analysing on the basis of SCM activity in comparison to our method of categorising trials on the basis of RT.

## 4.0 Discussion

EMG activity of orbicularis oculi (OOc) and SCM are the most commonly used indicators of the presence of a startle response, being among the last to habituate (Carlsen et al., 2007; Castellote et al., 2017; Kofler, Müller, & Valls-Solè, 2006). However, OOc responses are characterised by an early-onset component (the eye-protective auditory or somatosensory blink reflex) which is more resistant to habituation and takes a separate route to the brainstem as opposed to the later occurring startle component, which is more amenable to habituation and is associated with the generalised skeletomotor response to startle (Brown et al., 1991; Valls-Solé, Kumru, & Kofler, 2008). It is difficult to distinguish the acoustic/somatosensory eyeblink response from the startle response in OOc EMG records (Brown et al., 1991), and as such, SCM has been argued to provide a key indication of the presence of a “true” startle. On the basis of the assumption that startle activity is a necessary condition for the StartReact effect, the presence of SCM activity when prepared movements are triggered by an intense sensory stimulus has thus been used in the literature to make inferences about the potential mechanisms underlying the StartReact effect which may rely on activation of startle circuits (Carlsen et al., 2007; Valls-Solé et al., 1999). Movements made in response to the intense stimulus which occur without measurable surface EMG activity in SCM have therefore been deemed to be voluntarily initiated movements and unrepresentative of the StartReact effect. Analysis of data on the basis of SCM activity has traditionally examined the difference between SCM+ trials and SCM− trials to determine what types of movements are amenable to StartReact and those which are not. When a statistically significant difference cannot be found between SCM+ and SCM− trials for a particular muscle or task, that particular muscle or task is deemed to be unamenable to StartReact (Carlsen, Chua, Inglis, Sanderson, & Franks, 2009; Carlsen et al., 2007; Honeycutt et al., 2013). Differences in the neurophysiological efferent connectivity between muscles which are or are not amenable to StartReact in accordance with this method of analysis are then used to assert the involvement of different brain regions in the StartReact effect. Our analyses suggest a flaw in this interpretation of data. Firstly, analysis of probe trials failed to confirm that RT data are bimodally distributed, which may be expected if triggering differs for StartReact versus volitional movements. Furthermore, when percentiles within a CDF are approximated to response times on trials with and without SCM activity, and RTs are split into these SCM+ and SCM− percentiles, for some movement types a large proportion of responses with long RTs which would otherwise be considered to be indicative of slower, voluntarily triggered responses, can be seen to occur in the presence of SCM activity. A number of SCM− responses are also present in the group of responses with shorter latencies that are equivalent in terms of RT to responses otherwise recognised as typical StartReact triggered movements. While some of these short latency movements may have been anticipatory, or SCM activity may have gone undetected by surface EMG, this finding along with the presence of SCM+ responses in late RTs clearly demonstrates that SCM activity is neither always necessary, nor always sufficient, to identify the short response times which are a hallmark of StartReact movements (Marinovic & Tresilian, 2016). While SCM activity tends to be more prominent for the shortest latency movements, this is likely a product of SCM activation being more probable when levels of motor preparation are high (Leow et al., 2018; MacKinnon, Allen, Shiratori, & Rogers, 2013; Marinovic & Tresilian, 2016). Therefore, SCM activity may not be a product of the engagement of a unique triggering circuit, but rather a by-product, along with short response latency, of elevated preparatory activity.

Examination of our Bayesian tests of association (Appendix D) may provide a means to interpret why differences are observed between SCM+ and SCM− responses for some tasks, and not for others. A statistically significant difference was observed between SCM+ and SCM− trials for Castellote and Kofler’s (2018) BB latency in the flexion task, BB latency in the combined pinch-flex task, and FDI latency in the combined pinch-flex task, but not for FDI latency in the pinch task. Similarly, for BB latency in the flexion task and combined pinch-flex task, and FDI latency in the combined pinch-flex task, our Bayesian test of association provided decisive evidence (Jeffreys, 1961) to support the dependence of SCM activity and percentile categorisation – indicating the presence of SCM activity was most often found for responses within the fastest percentiles of RT. However, for the pinch-only task, the Bayesian test of association provided weaker evidence to support the dependence of SCM activity on percentile categorisation. This suggests that for this task, SCM activity was not significantly more likely to occur with responses which had the fastest RTs and could occur across both short and long latency movements. This finding of weak evidence to support the dependence of SCM activity and percentile categorisation holds true for all muscles and tasks we have analysed which failed to indicate a statistically significant difference between SCM+ and SCM− responses (see Appendix B and Appendix D). We may therefore conclude that this difference depends on the distribution of SCM+ responses across the spectrum of RTs.

It has been previously proposed that a lack of RT difference between SCM+ and SCM− trials indicates that the StartReact effect could not be elicited in a certain muscle or task (Carlsen et al., 2009, 2007; Honeycutt et al., 2013). Our analyses here suggest that the failure to find a statistically significant RT difference between SCM+ and SCM− responses for a given response type does not indicate a specific mechanism has failed to be activated by the intense stimulus, but rather, a larger proportion of SCM+ responses at late RTs is likely to have obscured this difference. Therefore, the presence of SCM activity is an unreliable method to indicate whether a distinct StartReact mechanism which produces the shortest response latencies has been activated; regardless of whether this pathway acts through the bypassing of cortical circuits or through an engagement of a larger and more functionally relevant brain network. Making inferences about the underlying circuitry of StartReact responses is therefore likely to be unreliable when using surface EMG activity in SCM as a sole criterion for the classification of responses. Furthermore, studies which classify responses on the basis of SCM activity are prone to the loss of large amounts of data. For example, when SCM activity is required to classify responses, participants for whom no measurable SCM activity can be consistently observed must be excluded entirely from analyses. This leads to a reduction of statistical power, unnecessary burden to the participant, and the loss of time and resources. On the basis of the unreliability of SCM activity as a criterion to determine the triggering mechanisms of responses and the loss of data associated with using this neurophysiological indicator, we therefore propose the mechanisms underlying the StartReact effect may be further examined when responses are categorised via their latency.

We deemed responses at or below the 45^th^ percentile to be representative of responses at the shortest latencies which most often occur with SCM activity, and subsequently categorised responses at the 45^th^ percentile of RT or earlier into our fast onset response category for analysis. Those at the 55^th^ percentile or later were similarly categorised into our slower onset response category for analysis, representative of voluntarily triggered responses. Our analysis of a representative dataset (Castellote & Kofler, 2018) showed a significant interaction of percentile categorisation with task type, indicating responses from the target muscles may have differed depending on the task that they were engaged in. Examination of our extended analyses (Appendix E) however, does not consistently show this interaction of percentile categorisation with task type or muscle type across datasets, even in muscles which are thought to strongly differ in their efferent connectivity to subcortical brain areas (e.g. Marinovic et al., 2014). This may warrant further examination of a modulated benefit of the intense sensory probe on the triggering of movements which differ in their neurophysiological connectivity. Furthermore, our Bayesian test of association analyses presented in Appendix D suggests there may be task-related factors that influence the dependence of SCM activity on RT. The percentile-SCM dependence that we observed for some tasks but not for others may be a consequence of high levels of motor preparation, however the task specificity we observed in these analyses may also suggest that it is possible for differences in the circuitry used for motor control to influence the distribution of SCM activity across RTs and as such, influence interpretations of the presence of the StartReact phenomenon. Such task-specific effects may relate to the use of SCM as part of a proximal stabilisation pattern in startle. Potentially, SCM may be activated to stabilise the body before rapid muscle activity in a proximal effector. This pattern of stabilisation may not be required as prominently for rapid activity in a more distal effector, which may provide some explanation for why the RT-SCM dependence was weaker for Castellote and Kofler’s (2018) distal pinch-only task, but was decisive for the remaining tasks which recruited the proximal BB.

## 5.0 Conclusions

Overall, inferences made about the presence of a distinct triggering mechanism for StartReact responses based on the presence or absence of SCM activity require careful consideration. The findings here suggest there are task- and muscle-specific responses to the probe stimulus that may influence both the manifestation of the StartReact as well as the ability to detect StartReact on the basis of SCM activity. Furthermore, while our analyses here cannot confirm nor rule out distinct triggering mechanisms for prepared motor responses via intense sensory stimuli, we suggest these underlying mechanisms for the StartReact effect should be further examined on the basis of response latency, rather than surface EMG activity of the SCM alone.

## Abbreviations

AD: anterior deltoid
APB: abductor pollicis brevis
BB: biceps brachii
Br: brachioradialis
CDF: cumulative distribution function
EMG: electromyogram
EDC: extensor digitorum communis
FDI: first dorsal interosseous
FDS: flexor digitorum superficialis
OO: orbicularis oris
OOc: orbicularis oculi
PD: posterior deltoid
Pe: pectoralis
RT: reaction time
SCM: sternocleidomastoid
TB: triceps brachii

## Appendix A

We conducted Hartigan’s (1985) dip test to test the multimodality of all datasets. The test failed to reject the null hypothesis of unimodality for all datasets we analysed. This suggests responses to intense sensory stimuli tend to fit a unimodal distribution. Resulting *p* values of the tests are reported in Table A.1.

**Table A.1.**
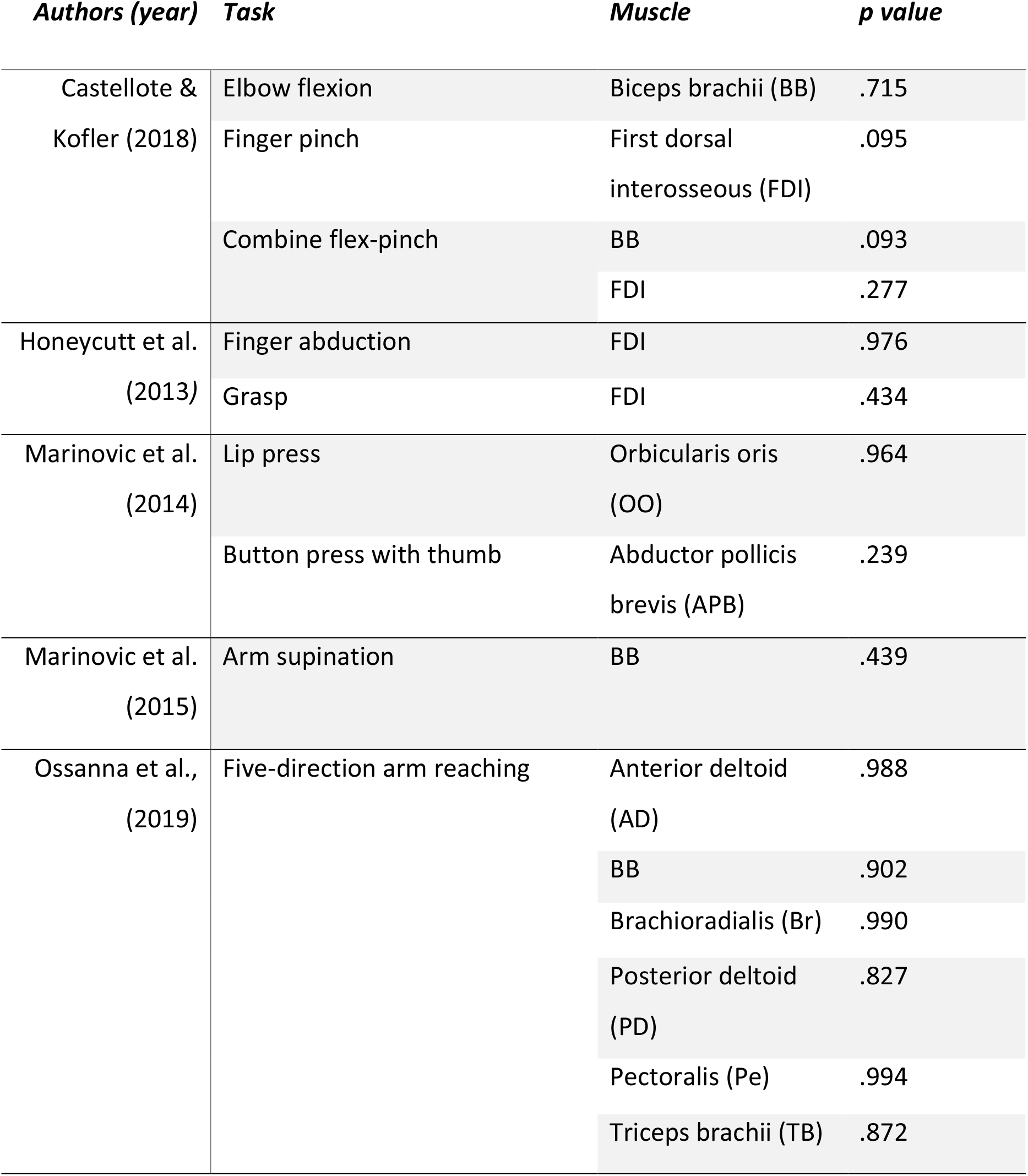
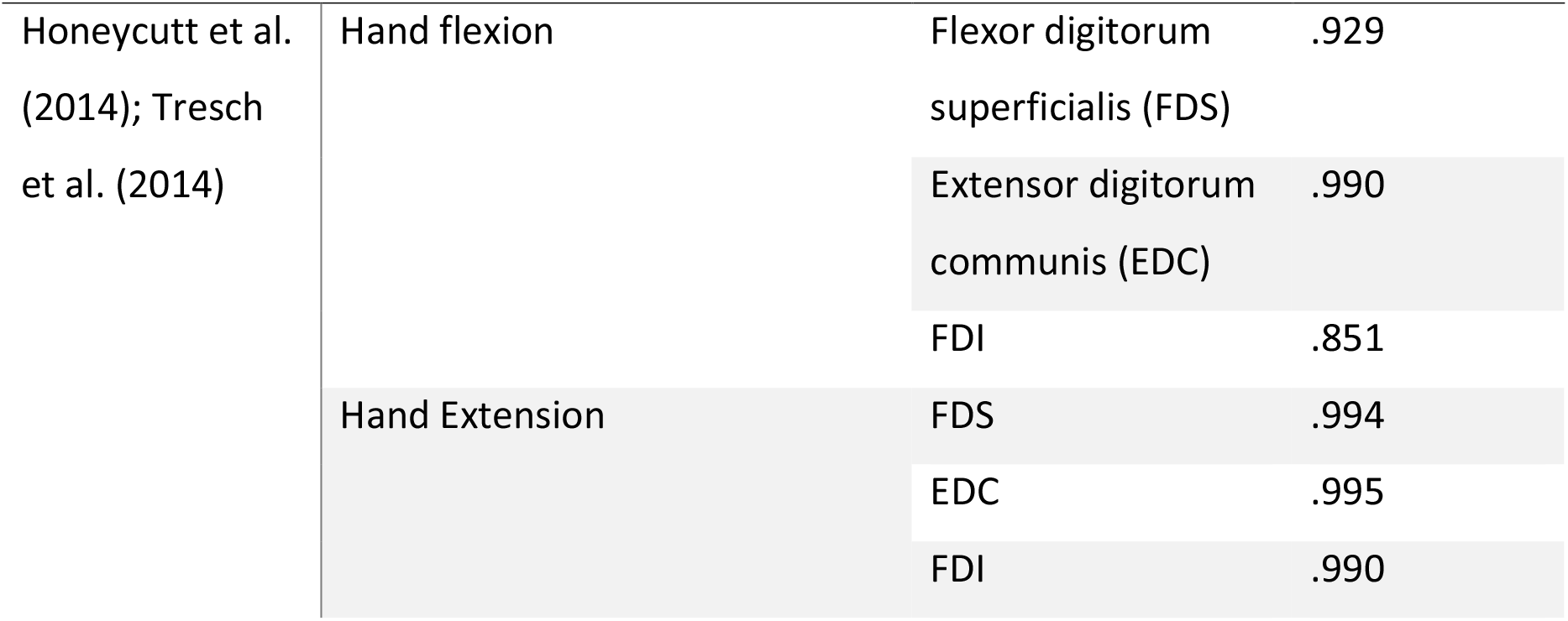
**values returned from our Hartigan’s (1985) dip test.** We tested the null hypothesis of unimodality for each muscle/task. Statistical significance is determined at α = 0.05.

## Appendix B

We conducted paired samples t-tests of each subject’s median SCM+ and SCM− trial RT for each movement type across datasets to examine the difference in response latency between SCM+ responses and SCM− resonses. Mean differences and confidence intervals for each movement type are reported in Table B.1.

**Table B.1.**
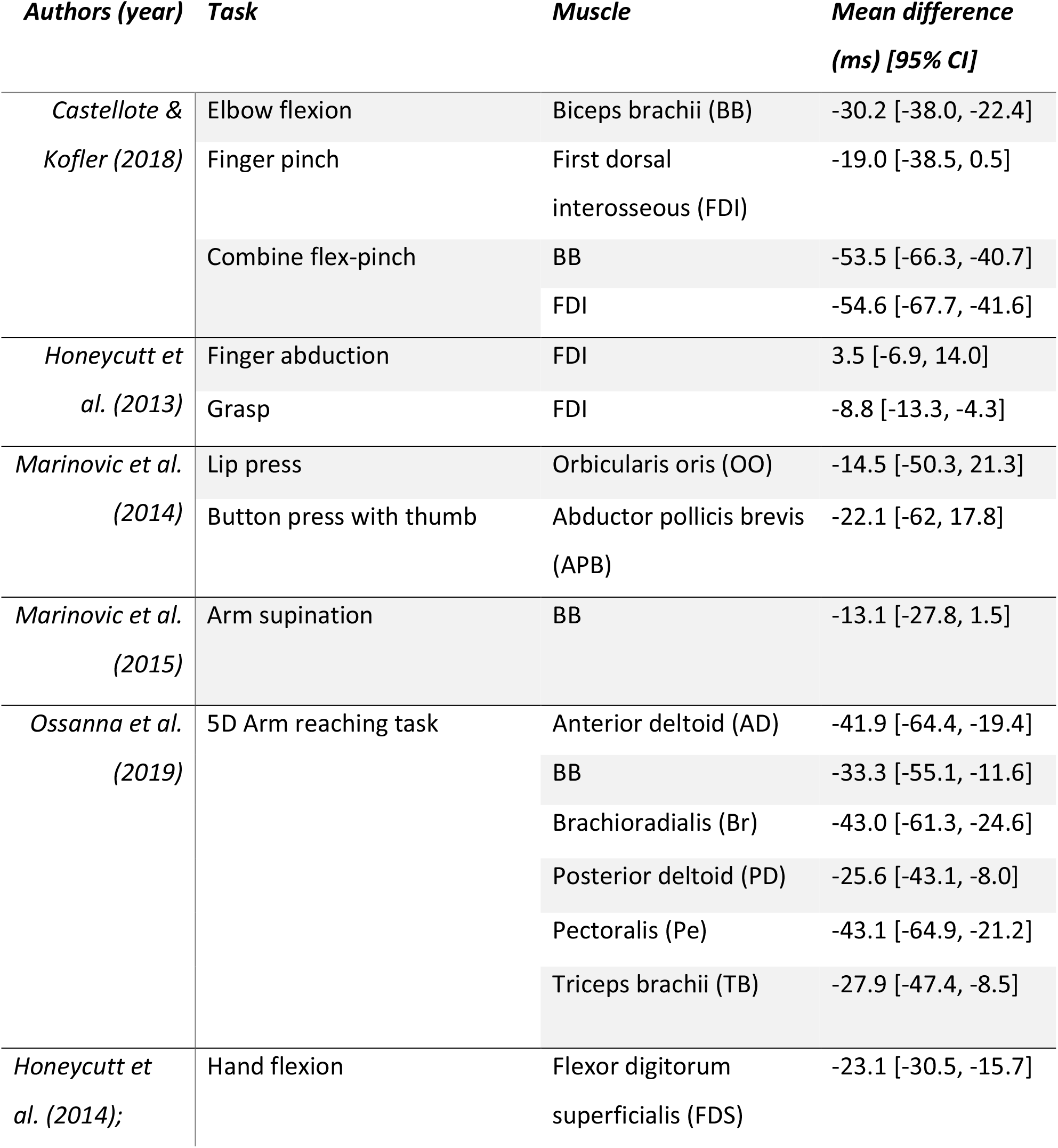
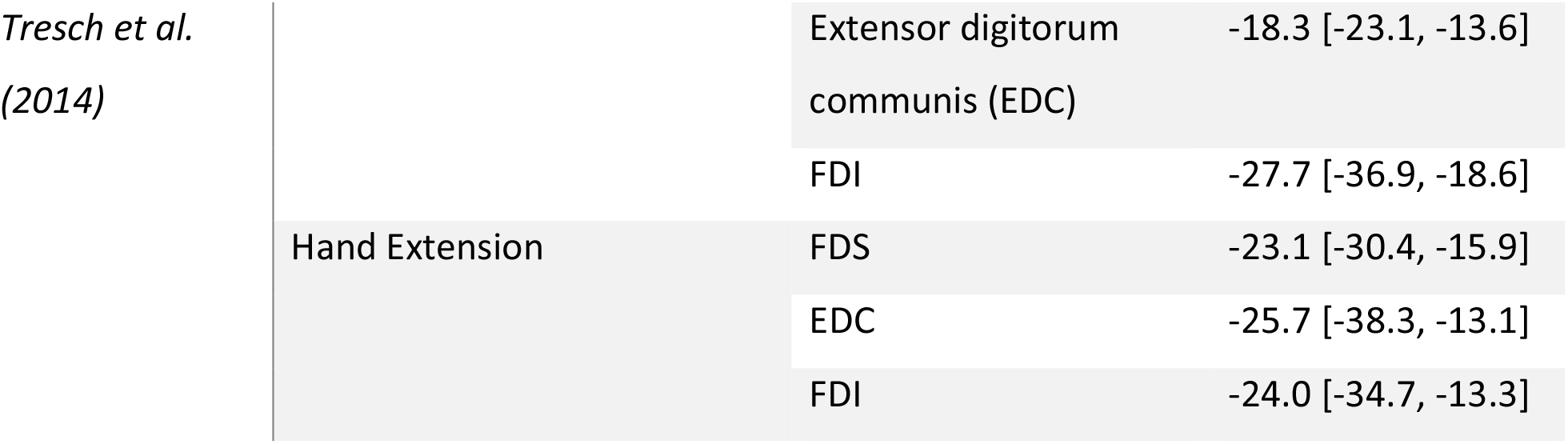
Difference between SCM+ and SCM− trials for each task and muscle analysed.

## Appendix C

We calculated CDFs for each muscle and task across all datasets analysed. For each movement type, the mean latency across participants of the percentile that closest matched the mean latency of SCM+ responses was deemed the SCM+ percentile for that task. Similarly, the SCM− percentile was determined as the percentile within the CDF that closest matched the mean latency of SCM− responses for that task. The CDFs for each movement type are plotted along with the mean latency of SCM+ and SCM− responses in Figures C.1 – C.5. Responses were placed into Startle and Non-Startle categories based on SCM+ and SCM− percentile latency (see Figure 1), and we subsequently calculated the percentage of responses within each category that occurred with SCM activity to determine the distribution of SCM+ responses between our categories. The distribution of SCM+ responses within the Startle and Non-Startle categories is displayed in Table 2 and Figures C.6 – C.10.

**Figure C.1.**
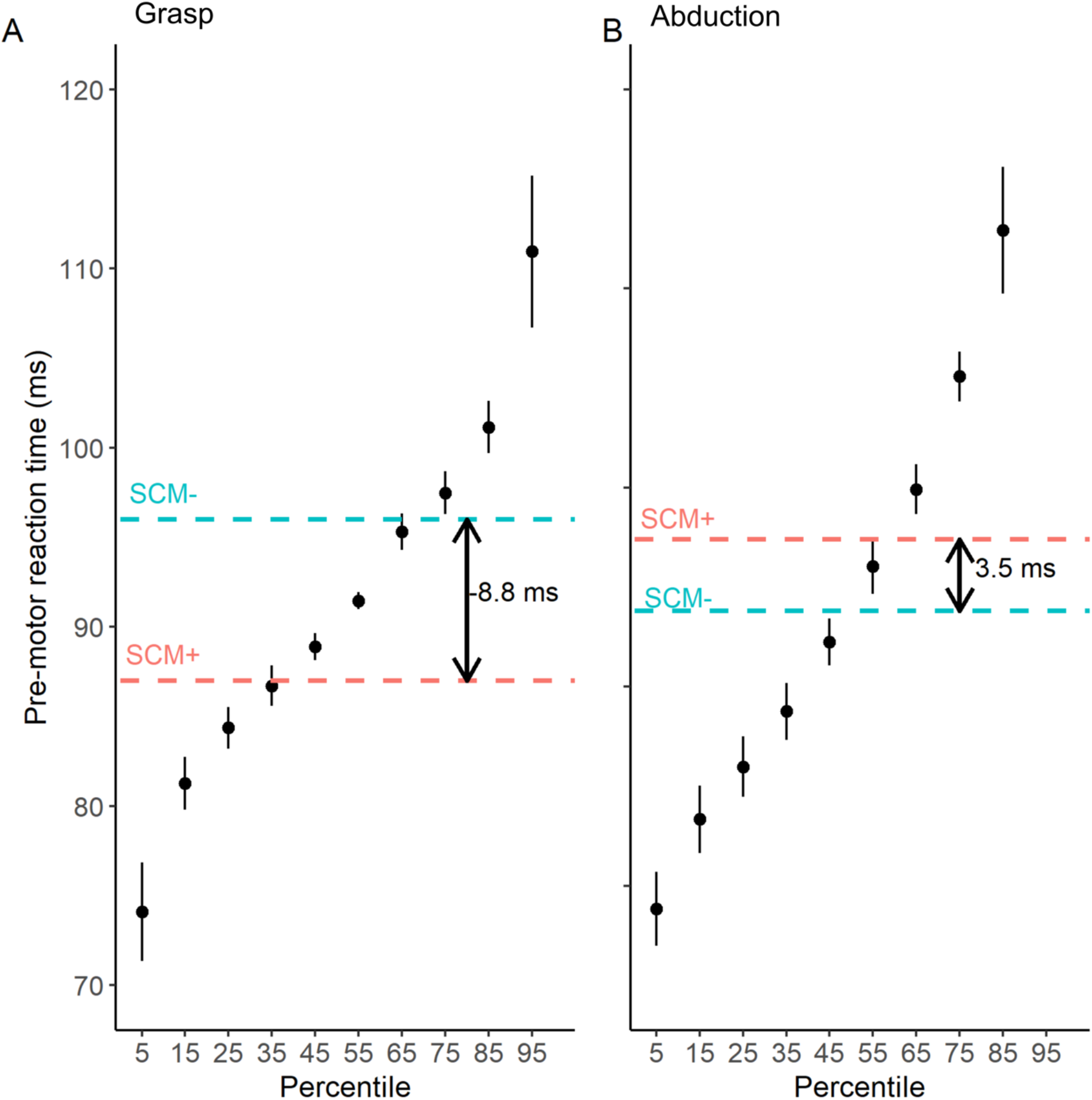
**Cumulative distribution function of first dorsal interosseous latency in Honeycutt et al.’s (2013) data.** A). Mean first dorsal interosseous (FDI) latency at each percentile of grasp task. B). Mean FDI latency at each percentile of finger abduction task. Mean latency of responses in the presence (SCM+) and absence (SCM−) of sternocleidomastoid activity are shown by the dotted lines. Error bars represent standard error of the mean.

**Figure C.2.**
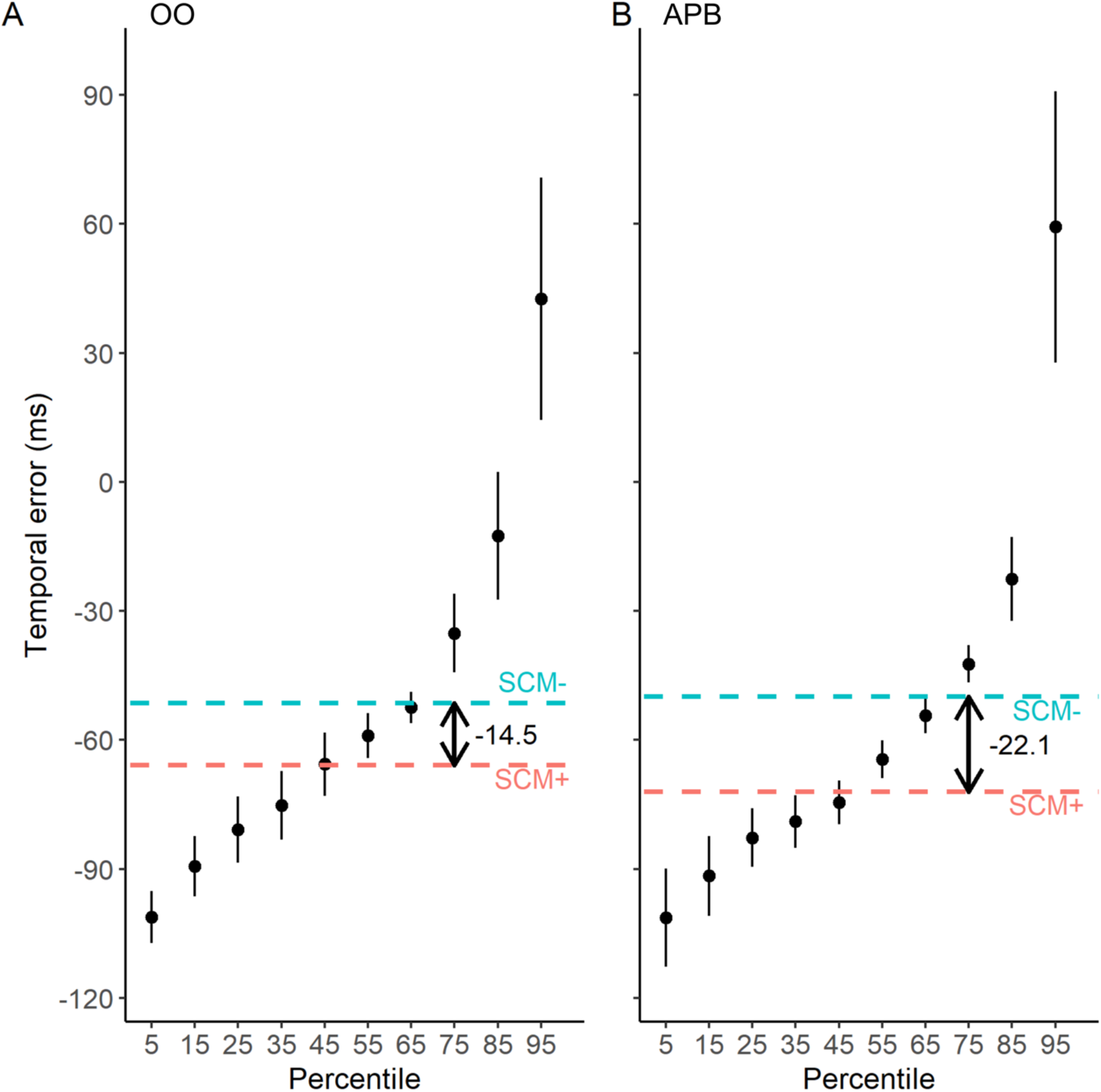
**Cumulative distribution function of temporal error in Marinovic et al.’s (2014) anticipatory timing tasks.** A). Mean temporal error of orbicularis oris (OO) at each percentile B). Mean temporal error of abductor pollicis brevis (APB) at each percentile. Mean latency of responses in the presence (SCM+) and absence (SCM−) of sternocleidomastoid activity are shown by the dotted lines. Error bars represent standard error of the mean.

**Figure C.3.**
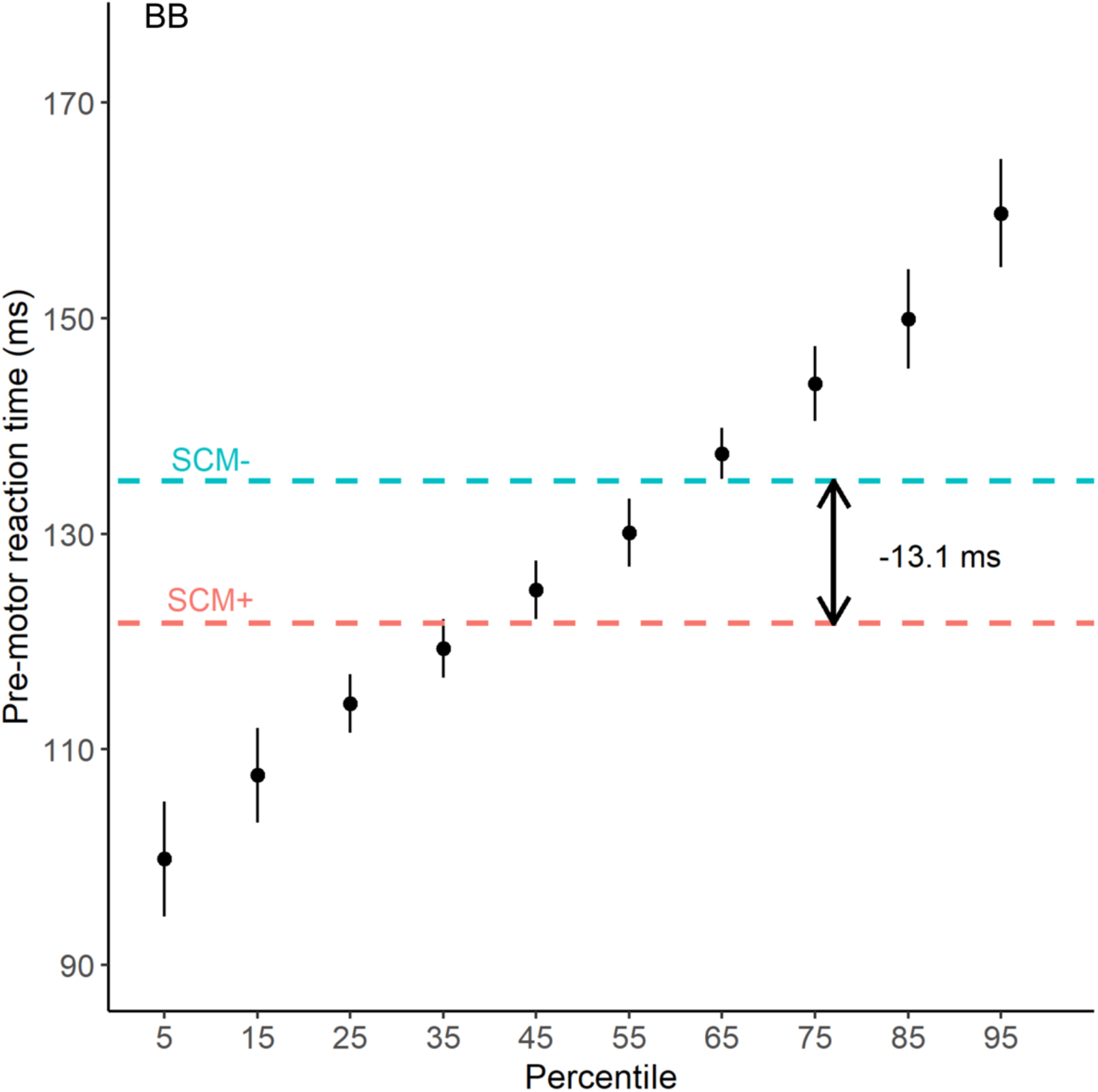
**Cumulative distribution function of biceps brachii latency in Marinovic et al.’s (2015) arm supination task.** Mean response times of biceps brachii (BB) at each percentile are displayed. Mean latency of responses in the presence (SCM+) and absence (SCM−) of sternocleidomastoid activity are shown by the dotted lines. Error bars represent standard error of the mean.

**Figure C.4.**
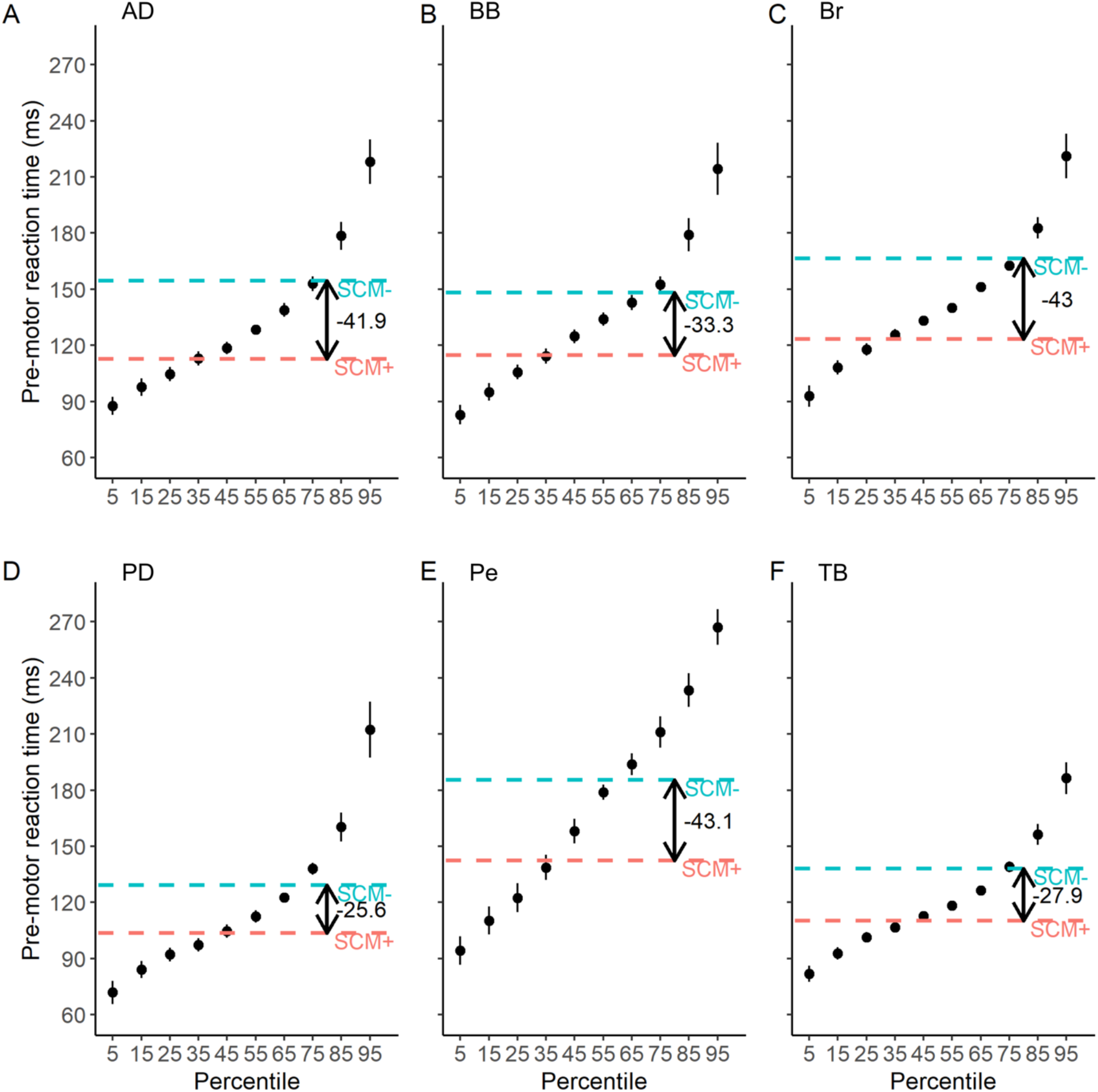
**Cumulative distribution function of response latencies in Ossanna et al.’s (2019) reaching task.** A). Mean latency of anterior deltoid (AD) at each percentile B). Mean latency of biceps brachii (BB) at each percentile C). Mean latency of Brachioradialis (Br) at each percentile D). Mean latency of posterior deltoid (PD) at each percentile E). Mean latency of pectoralis (Pe) at each percentile F). Mean latency of triceps brachii (TB) at each percentile. Mean latency of responses in the presence (SCM+) and absence (SCM−) of sternocleidomastoid activity are shown by the dotted lines. Error bars represent standard error of the mean.

**Figure C.5.**
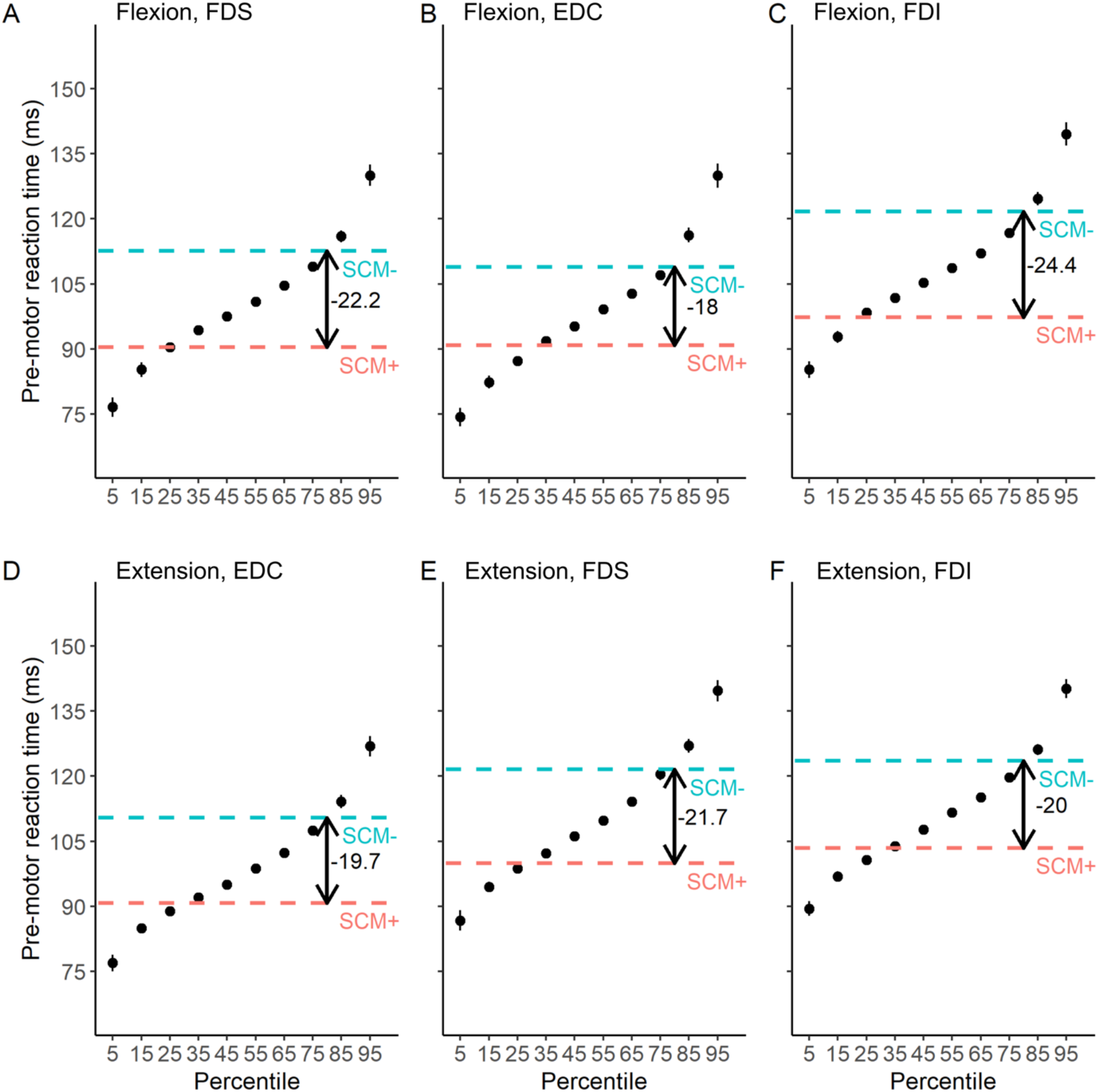
**Cumulative distribution function of response latencies in Honeycutt et al. (2014); Tresch et al. (2015).** A). Mean latency of flexor digitorum superficialis (FDS) at each percentile of the hand flexion task. B). Mean latency of extensor digitorum communis (EDC) at each percentile of the hand flexion task. C). Mean latency of first dorsal interosseous (FDI) at each percentile of the hand flexion task. D). Mean latency of EDC at each percentile of the hand extension task. E). Mean latency of FDS latency at each percentile of the hand extension task. F). Mean latency of FDI at each percentile of the hand extension task. Mean latency of responses in the presence (SCM+) and absence (SCM−) of sternocleidomastoid activity are shown by the dotted lines. Error bars represent standard error of the mean.

**Figure C.6.**
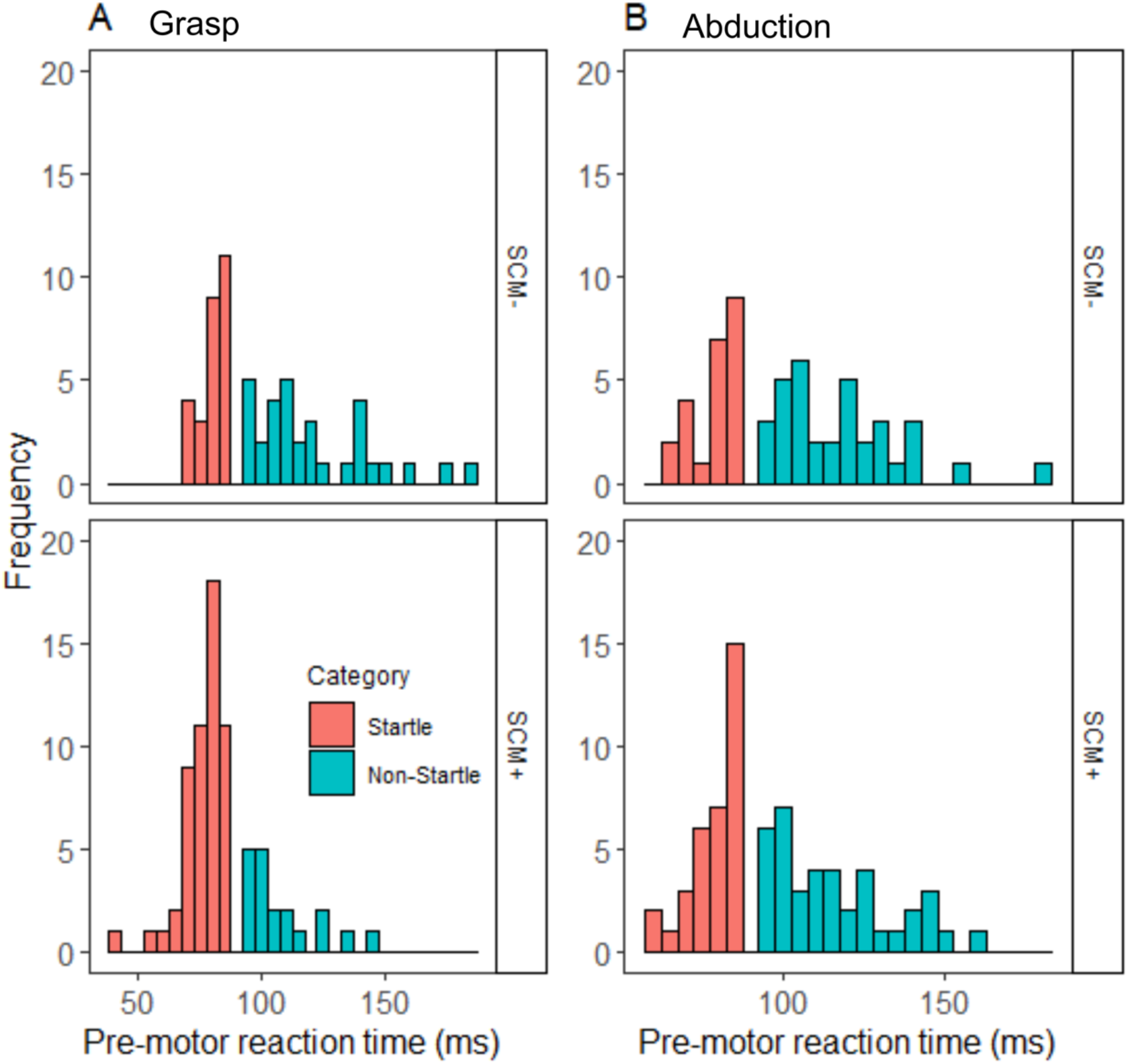
**Histogram displaying frequency of response times in Honeycutt et al.’s (2013) data.** SCM+ and SCM− responses across Startle and Non-Startle percentile categories are displayed. A). First dorsal interosseous (FDI) latency in grasp task. B). FDI latency in finger abduction task.

**Figure C.7.**
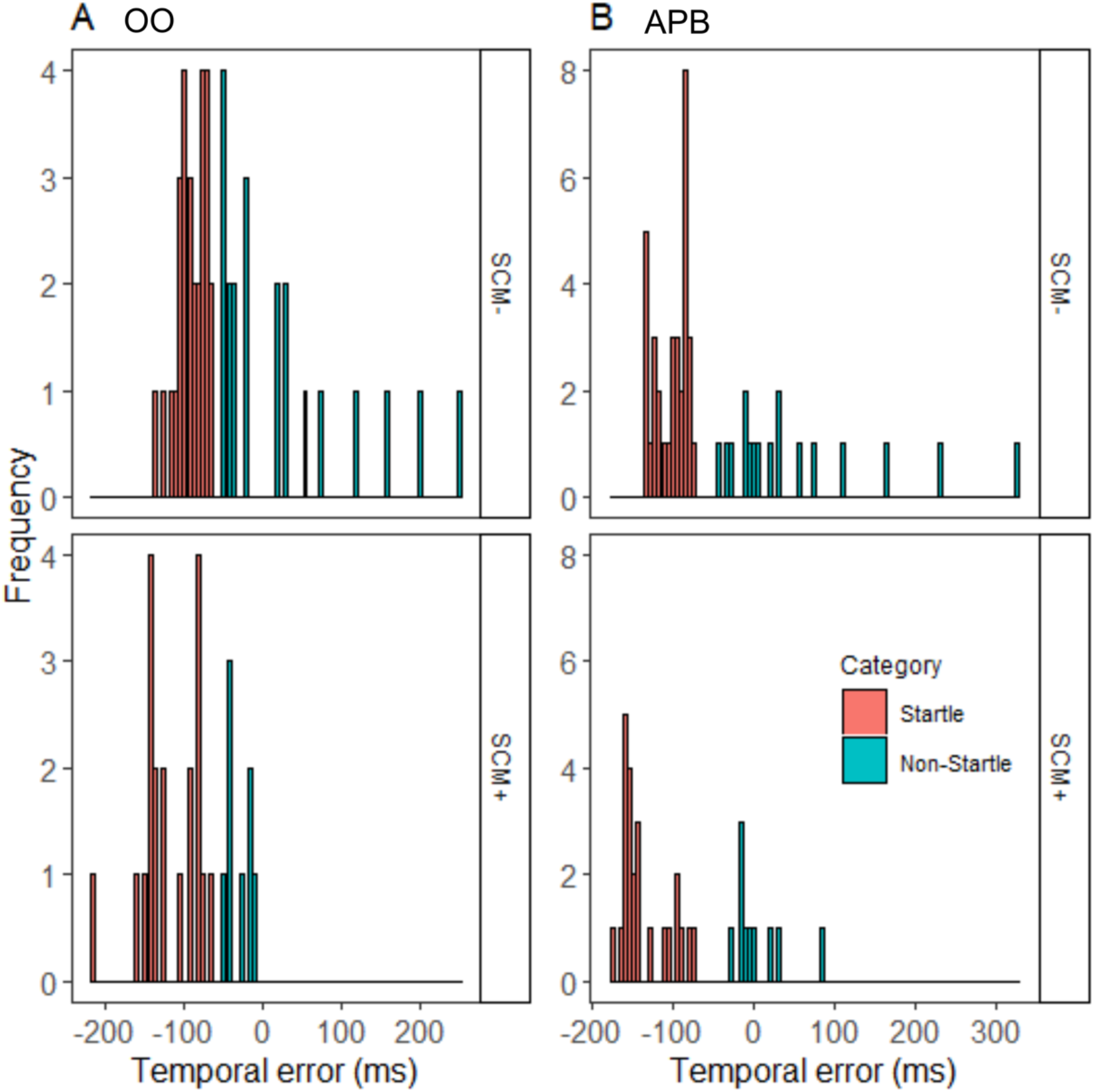
**Histogram displaying frequency of temporal error in Marinovic et al.’s (2014) data.** SCM+ and SCM− responses across Startle and Non-Startle percentile categories are displayed. A). Temporal error of orbicularis oris (OO). B). Temporal error of abductor pollicis brevis (APB).

**Figure C.8.**
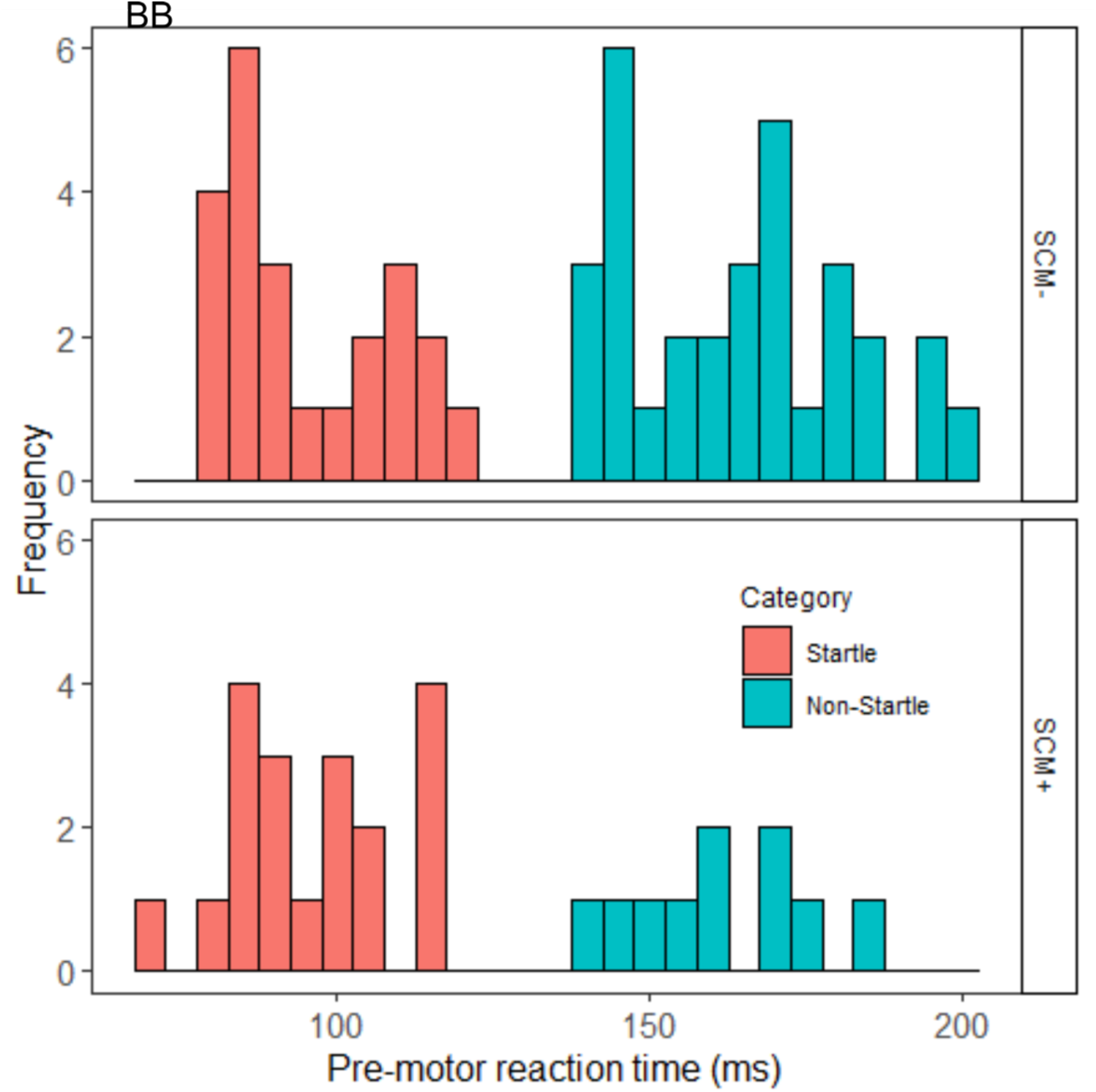
**Histogram displaying frequency of response times of biceps brachii in Marinovic et al.’s (2015) data.** SCM+ and SCM− responses across Startle and Non-Startle percentile categories are displayed. BB = biceps brachii.

**Figure C.9.**
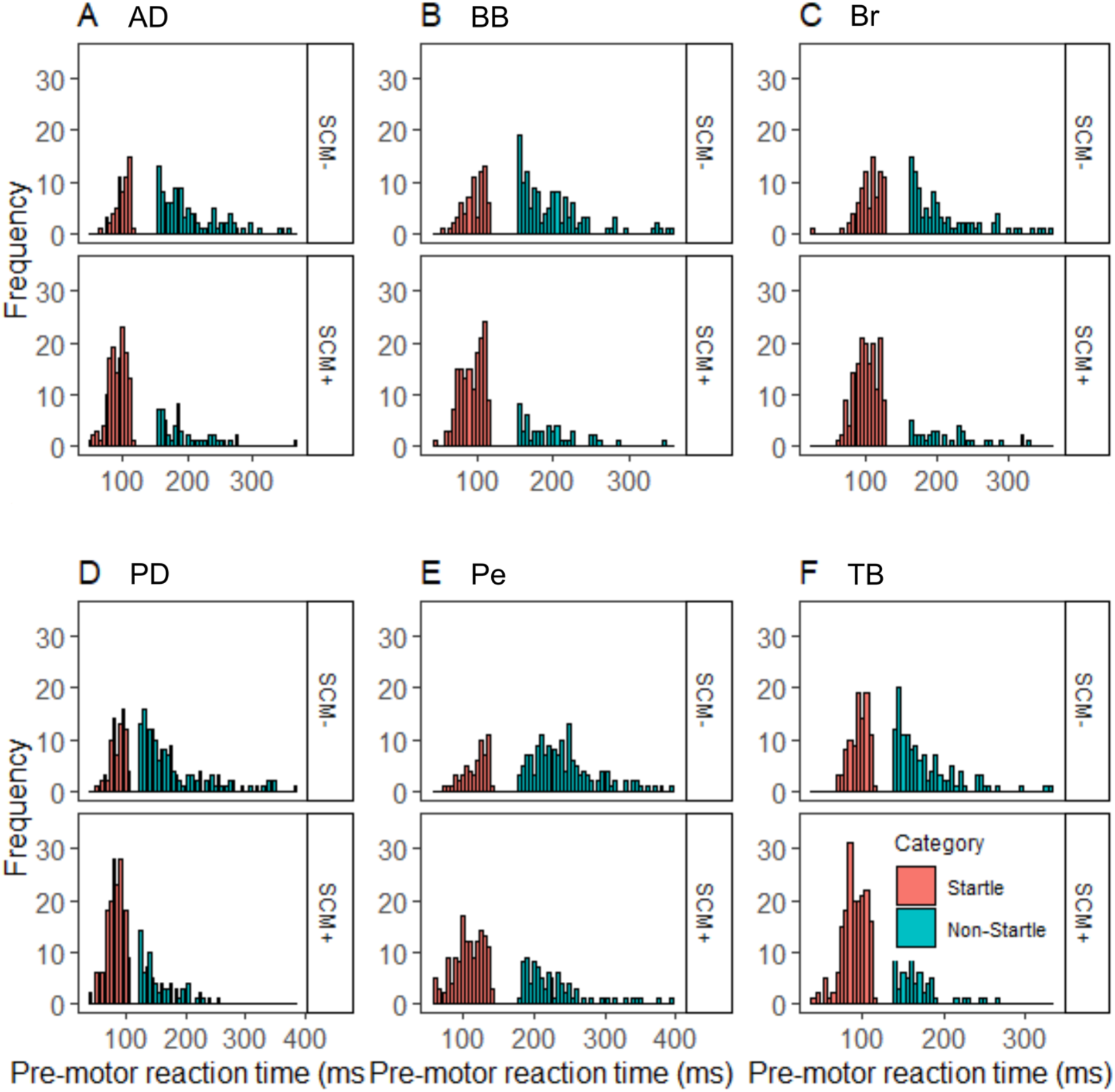
**Histogram displaying frequency of response times in Ossanna et al.’s (2019) data.** SCM+ and SCM− responses across Startle and Non-Startle percentile categories are displayed. A). Anterior deltoid (AD) latency B). Biceps brachii (BB) latency C). Brachioradialis (Br) latency D). Posterior deltoid (PD) latency E). Pectoralis (Pe) latency F). Triceps brachii (TB) latency.

**Figure C.10.**
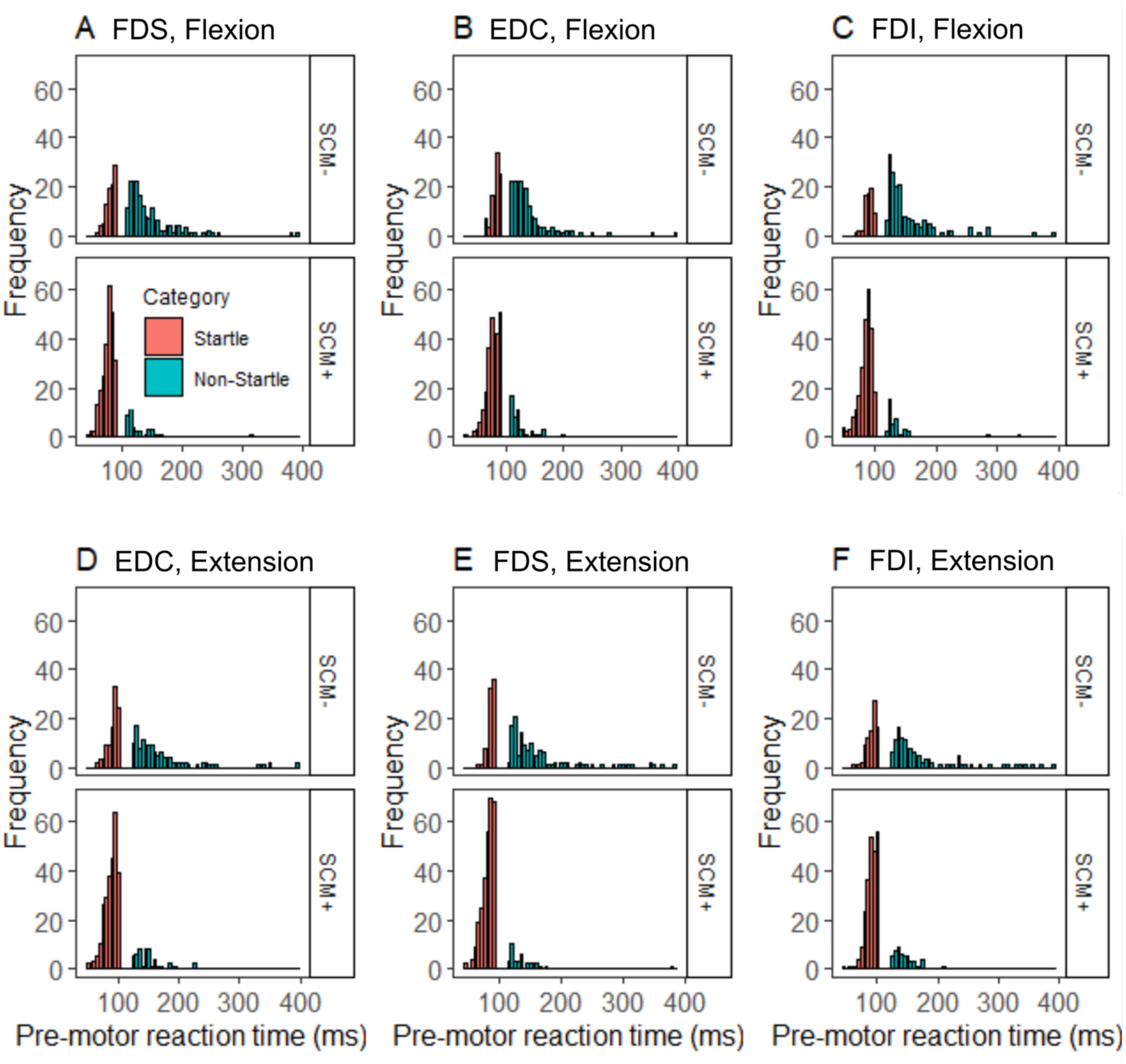
**Histogram displaying frequency of response times in Honeycutt et al. (2014) and Tresch et al.’s (2014) data.** SCM+ and SCM− responses across Startle and Non-Startle percentile categories are displayed. A). Flexion task, flexor digitorum superficialis (FDS) latency. B). Flexion task, extensor digitorum communis (EDC) latency. C). Flexion task, FDI latency. D). Extension task, EDC latency. E). Extension task, FDS latency. F). Extension task, FDI latency.

## Appendix D

Given a large proportion (max = 56.32%) of responses in the Non-Startle categorisation of RT occurred with SCM activity (see Table 2), we conducted a series of Bayesian tests of association (Albert, 1997) to examine whether the presence of SCM activity depends on our Startle and Non-Startle RT categories. The resulting Bayes Factors (BFs) are reported in Table D.1.

**Table D.1.**
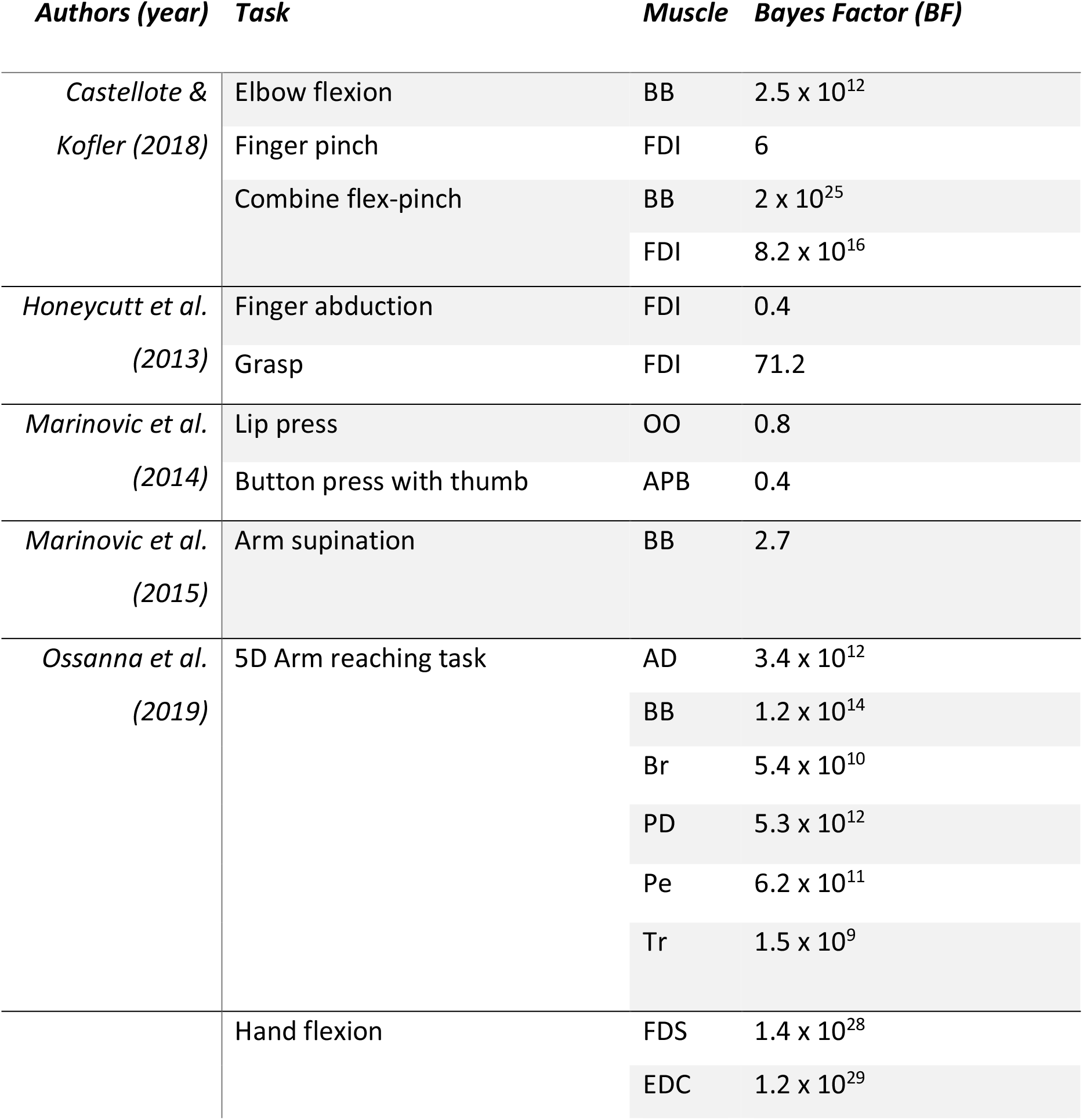
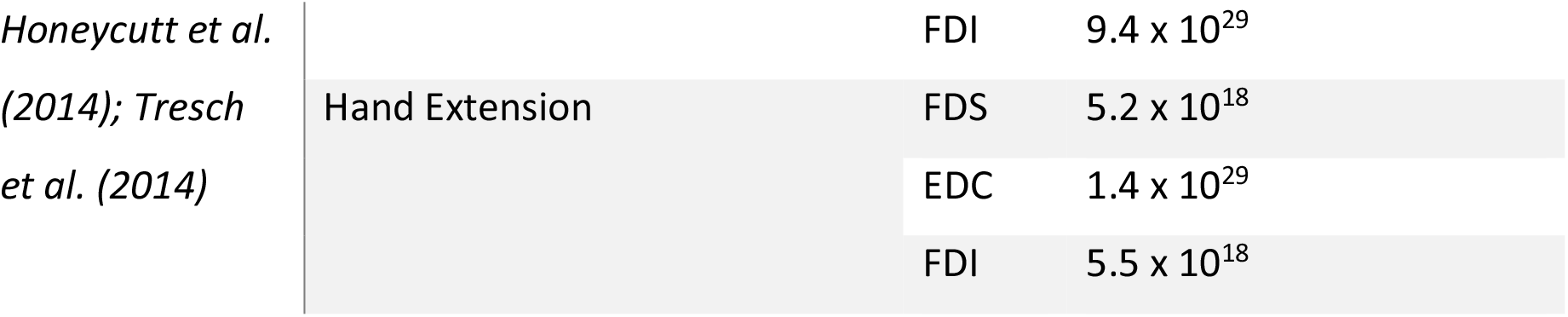
Bayes factors calculated by Bayesian tests of association for each movement type across all datasets analysed. Bayes Factors (BFs) indicate the degree of evidence to support the dependence of SCM activity and response latency categorisation. BF = 1 indicates no support for the null or alternative hypothesis, BF > 3 indicates substantial evidence, BF > 10 indicates strong evidence, BF > 30 indicates very strong evidence, and BF > 100 indicates decisive evidence for the alternative hypothesis (Jeffreys, 1961).

## Appendix E

Our analyses provided weak evidence to support the hypothesis that the presence of SCM activity is always dependent on percentile categorisation. That is, a significant proportion of SCM+ responses are not only found in the Startle category, but also within the Non-Startle category which approximates SCM− response latencies. Therefore, we examined an alternative approach for investigating triggering mechanisms of responses via intense sensory stimuli: categorisation via percentiles of RT. Response times at the 45^th^ percentile or earlier – the fast onset percentiles - were likely to be indicative of responses which occur more often in the presence of SCM activity and which are likely to be indicative of any distinct neurophysiological mechanism responsible for the StartReact effect that may be present. Similarly, response times at the 55^th^ percentile or later were chosen to represent the slower onset responses which less frequently occur with SCM activity. We conducted a series of linear mixed-effects models on each dataset using these percentile categories (means shown in Figure E.1) to examine any interactions of percentile categorisation with the muscle and task factors to determine whether differing neurophysiological contributions to different movement types alter their benefit received from the intense auditory probe. The resulting statistics are shown in Table E.1.

**Table E.1.** Statistical output of our linear mixed effects models.

| *Authors*<br>*(year)* | *Main Effect/Interaction* | *df* | *F* | *p* |
| --- | --- | --- | --- | --- |
| *Castellote & Kofler*<br>*(2018)* | Percentile | 1, 422 | 533.7 | <.001 |
|  | Task | 1, 422 | 32.5 | <.001 |
|  | Muscle | 1, 422 | 26.6 | <.001 |
|  | Percentile*Task | 1, 422 | 18.6 | <.001 |
|  | Percentile*Muscle | 1, 422 | 0.0 | .814 |
| *Honeycutt et al.*<br>*(2013)* | Percentile | 1, 187 | 158.1 | <.001 |
|  | Task | 1,187 | 11.1 | .001 |
|  | Percentile*Task | 1, 187 | 2.7 | .099 |
| *Marinovic et al.*<br>*(2014)* | Percentile | 1, 130 | 62.2 | <.001 |
|  | Muscle | 1, 130 | 0.1 | .745 |
|  | Percentile*Muscle | 1, 130 | 0.0 | .905 |
| *Marinovic et al. (2015)* | Percentile | 1, 89 | 103.7 | <.001 |
| *Ossanna et al. (2019)* | Percentile | 1, 579 | 501.8 | <.001 |
|  | Muscle | 5, 579 | 29.9 | <.001 |
|  | Percentile*Muscle | 5, 579 | 5.2 | <.001 |
| *Honeycutt et al. (2014); Tresch et al. (2014)* | Percentile | 1, 1944 | 825.7 | <.001 |
|  | Task | 1, 1944 | 43.8 | <.001 |
|  | Muscle | 2, 1944 | 27.5 | <.001 |
|  | Percentile*Task | 1, 1944 | 2.6 | .108 |
|  | Percentile*Muscle | 2, 1944 | 0.1 | .899 |

**Figure E1.**
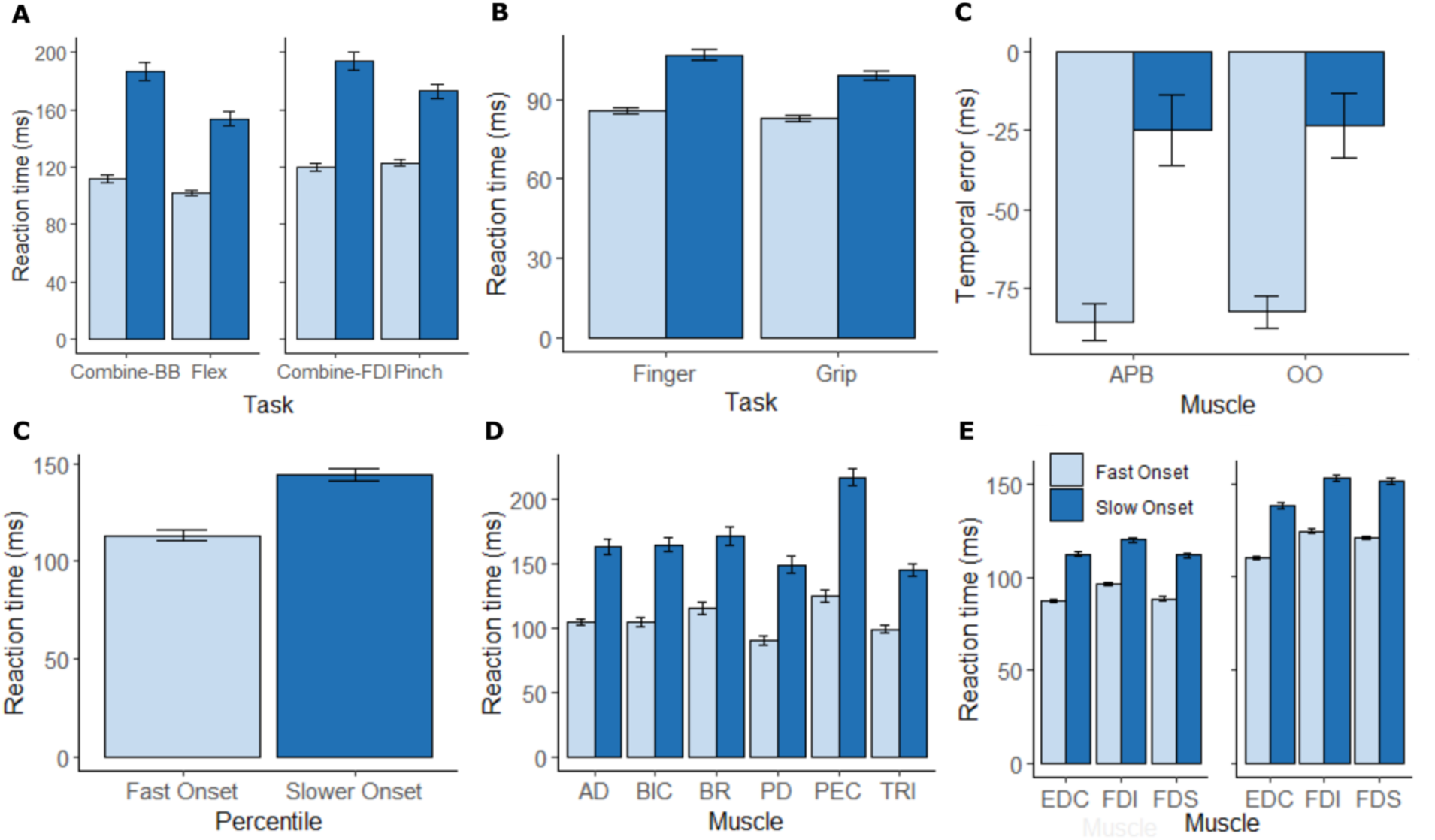
Mean response times across datasets. **A)** Mean RT for Fast and Slow Percentiles across muscles and tasks in Castellotte and Kofler (2018). **B)** Mean RT for Fast and Slow Percentiles in Honeycutt et al.’s (2013) finger and grip tasks. **C)** Mean temporal error for Fast and Slow Percentiles across muscles and tasks in Marinovic et al. (2014). **D)** Mean RT for Fast and Slow Percentiles Marinovic et al.’s (2014) arm supination task. **E)** Mean RT for Fast and Slow Percentiles across muscles recorded in Ossanna et al. (2019). **F)** Mean RT for Fast and Slow Percentiles across muscles and tasks in Honeycutt et al. (2014), Tresch et al. (2014). Error bars represent standard error of the mean.

